# RNA labelling in live plants reveals single cell transcriptional dynamics: application to phosphate signaling

**DOI:** 10.1101/2021.04.12.439441

**Authors:** Sahar Hani, Laura Cuyas, Pascale David, David Secco, James Whelan, Marie-Christine Thibaud, Rémy Merret, Florian Mueller, Nathalie Pochon, Hélène Javot, Orestis Faklaris, Eric Maréchal, Edouard Bertrand, Laurent Nussaume

## Abstract

Plants are sessile organisms constantly adapting to ambient fluctuations through spatial and temporal transcriptional responses. Here, we implemented the latest generation RNA imaging system and combined it with microfluidics to visualize transcriptional regulation in living Arabidopsis plants. This enabled quantitative measurements of the transcriptional activity of single loci in single cells, real time and changing environmental conditions. Using phosphate responsive genes as model, we found that active genes displayed high transcription initiation rates (∼3s) and frequently clustered together in endoreplicated cells. We observed gene bursting and large allelic differences in single cells, revealing that at steady-state, intrinsic noise dominated extrinsic variations. Moreover, we established that transcriptional repression triggered in roots by phosphate, a crucial macronutrient limiting plant development, occurred with unexpected fast kinetics (∼minutes) and striking heterogeneity between neighboring cells. Access to single cell RNA polymerase II dynamics within live plants will benefit future studies of signaling processes.

## Introduction

Plants are sessile organisms permanently coping with environmental variations. Transcriptional reprogramming^1^ plays a key role in these responses as illustrated by the number of plant transcription factors (5.4% in Arabidopsis, https://agris-knowledgebase.org/). Most physiological studies quantify mRNA abundance in organs. Accessing specific cell types is possible by expressing a fluorescent marker in cells of interest, to FACS sort them and perform sequencing analysis^2^. Nevertheless, this requires long enzymatic digestion to generate protoplasts, preventing fast kinetic studies. Transcriptional fusions between promoters and reporter genes such as GFP provide cellular resolution, but the time required to accumulate detectable level of mature fluorophore (often in the range of tens of minutes^3^) prevents the rapid transcriptional monitoring. This is moreover highly variable and depends on the strength of the promoter studied and the microscopy setup, leading to confounding effects. These considerations are even more crucial when studying transcriptional inhibition. The above-mentioned experiments provide access to total RNA (or protein), resulting from the balance between synthesis and degradation. Many fluorescent proteins decay in the range of hours and RNA degradation is highly variable from one gene to another^4,5^. For instance, the median half-life value of *Arabidopsis* mRNAs is 107 min, but exceeds one day for many messengers^4^. This is far too long for transcriptional responses occurring within seconds such as for instance during light stress ^6^.

All these issues can be resolved with a method originally developed for yeast and animal granting direct real time access to transcriptional activity at the level of single cells^7-10^. It is based on a fusion between GFP and the bacteriophage MS2 coat protein (MCP-GFP)^7,8^ or on related systems^11^. The MCP-GFP recognizes a specific RNA stem-loop inserted in multiple copies into a reporter RNA, promoting MCP-GFP multimerization to provide signal bright enough for single RNA visualization^9^. Importantly, binding occurs during RNA synthesis and monitors transcription in real time. In plants, this technology would offer major advantages for physiological studies such as adaptations to biotic and abiotic stresses. Indeed, it grants access to the variability of all cell types providing ways to understand how the activity of single cells is integrated within tissues or organs and thus to better understand gene regulation. Real time quantitative analysis can be performed for transcription initiation, elongation or gene bursting. This last phenomenon can further be used to identify promoter states that are rate-limiting for transcription initiation, and thus likely points of regulation^12^. The MS2 technology also allows the identification of extrinsic and intrinsic sources of transcriptional noise

Therefore, we implemented it into plants using state-of the art MS2×128 repeats to tag a single RNA with ∼250 to 500 GFPs for optimal detection sensitivity^13^. We used this system to analyze the transcriptional response to a major macronutrient: inorganic phosphate (Pi). Pi deficiency triggers major transcriptional modifications affecting plant development and metabolism ^14,15^. These are mainly controlled by master regulator genes of the PHR1 family^15-17^. Being constitutive, their regulatory activity relies on inhibitors of the SPX family tuned by Pi uptake^18,19^ as they inhibit PHR1 activity only in the presence of Pi metabolites ^20,21^.

Here we used the promoter of early Pi responding genes to drive transcription of an MS2×128 reporter. We combined fast quantitative imaging and microfluidics^22^, to precisely control Pi delivery while providing stable environmental conditions. It revealed a high heterogeneity of responses between adjacent cells and identified the rapid perception of Pi by the root (within 3/5 min) validating the power of the MS2 technology to dissect plant transcriptional regulation.

## Results

### Phosphate resupply promotes rapid transcriptional modifications

To identify early markers sensitive to phosphate, we starved plants and performed Pi refeeding experiments. After addition of Pi in the liquid culture medium, we harvested roots and leaves after 30, 60 and 180 min for RNA-seq analysis. In roots, only 22 genes exhibited a significant two-fold reduction of their transcript level over the three time points (Fig. S1A-B). Analysis of the shoot samples revealed a delayed reduction of all these markers in aerial parts (only one third were repressed after 30 min) indicating that Pi was first perceived by roots. Independent experiments analyzed by RTqPCR using well known markers for Pi starvation^15,23^, involved in Pi uptake (*PHT1;4*) or Pi-induced metabolic remodelling (*SQD2*), confirmed these results (Fig. 1A and S1B). The rapid down-regulation observed also highlighted the fast turn over of these transcripts (half-life estimated to 15-30 minutes; Fig. S1A-B), in regard to the Arabidopsis median value of 3.8 hours^5^. For the rest of this study we selected two genes (*SPX1* and At5G20790, named hereafter *UNI1* for Unicorn1), which combined the highest levels of expression in Pi depleted medium (-Pi) with broad dynamics as measured by +Pi/-Pi fold change.

**Figure 1:**
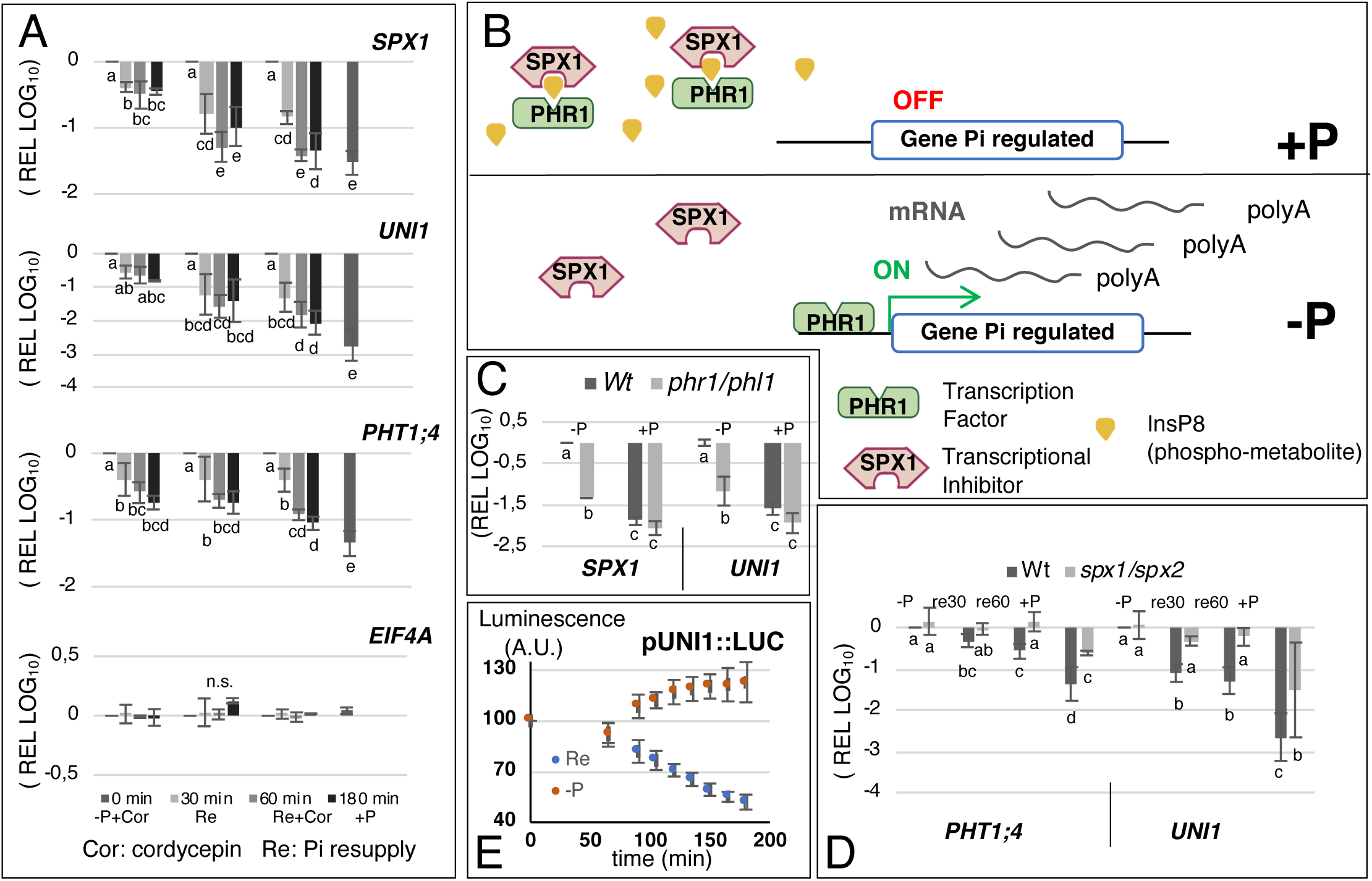
Identification of fast responsive transcripts regulated at transcriptional level by phosphate resupply. (A) RTqPCRs of roots of seedlings grown for 7 days either in the presence of Phosphate (+Pi), in the absence of Phosphate (-Pi), or in the absence of Phosphate for 7 days followed by Phosphate resupply for 30 minutes (Re30) or 60 minutes (Re60), or 180 minutes. *EIF4A,* a translation initiation factor known not to react to cordycepin addition was used as control^45^; n =2-3. (B) Model depicting the transcriptional regulation by phosphate. (C) RTqPCRs of roots of WT and *phr1/phl1* seedlings grown during 7 days under +Pi and –Pi; n=2-3. (D) RTqPCRs of roots of WT and *spx1/spx2* seedlings grown during 7 days under +Pi or -Pi, and supplemented with Pi for 30 or 60 minutes; n=3-5. (E) Kinetic of luminescence measurements in pUNI1::LUC transgenic seedlings between –Pi and resupplied sample, value are relative to –Pi; n=5. For all RTqPCR experiments *TUBULIN* was used as a housekeeping reference gene for normalization. Values are log_10_ relative expression levels (REL) normalized to 1 for -Pi levels at time zero represented. Different letters indicate significantly different means (SNK one-way ANOVA, p<0.05, Rstudio). Error bars represent standard deviation and n is the number of biological replicates used for RNA extraction.

Regulation was assumed to occur mainly transcriptionally, as the transcriptional inhibitor cordycepin mimicked effect of Pi resupply (Fig. 1A). This was confirmed by transcriptional fusions with luciferase, which conferred to the reporter gene a similar temporal response to Pi resupply (Fig. S1C), whereas fusing a CaMV 35S constitutive promoter to the coding region of the markers did not reveal any significant difference of RNA levels between +Pi (after 180 min refeeding) or -Pi conditions (Fig. S1D), except for short transient induction promoted by the stress of Pi addition observed during first hour following resupply. A sequence analysis of the 22 genes identified above revealed the presence of P1BS regulatory boxes^16^ in 95% of the cases. This box is the binding site of master regulators belonging to PHR1/PHL1 family. Addition of Pi promotes synthesis of inositol pyrophosphates, which act as a molecular tether to fix members of the SPX family to PHR1 and inhibit its activity^20,21^ (Fig. 1B). Consistently, the induction of *SPX1* and *UNI1* during Pi starvation was nearly fully abolished in *phr1/phl1* double mutants, which suppress a majority of PHR1/PHL proteins (Fig. 1C). Moreover, this was observed for 91% of the 22 genes identified (Fig. S1A^17^). SPX family members inhibit PHR1 and the analysis of *spx1/spx2* double mutants further revealed that reducing the SPX protein pool delayed the repression triggered by Pi addition (Fig. 1D). Altogether, these results demonstrated that the fast decrease of the 22 transcripts identified here resulted from transcriptional control involving several key players such as PHR or SPX family of proteins. Of note, classical transcriptional fusion with Luciferase did not report a fast transcriptional repression (Fig. 1E). More than one hour was required to observe a significant decrease of the signal, whereas RTqPCR failed to detect a significant reduction of mRNA levels for at least 15-20 minutes after resupply (data not shown). To better characterize the transcriptional response to Pi resupply, we implemented the MS2 system for RNA labeling, a technology so far restricted mostly to animal and yeast but offering unique spatio-temporal resolution to study transcriptional regulation.

### Generation and validation of MS2-tagged Arabidopsis lines

We used the last generation MS2 tag containing 128xMS2 repeats, originally developped to image single-molecule of HIV-1 RNA^13^. To improve RNA folding and prevent plasmid instability, this construct is made of 32 distinct MS2 stem loops replicated four times, with each stem-loop binding dimers of the MCP protein with sub-nanomolar affinity^13^ (Fig. 2A). In animal cells, this extended tag provides about a five-fold improved sensitivity over the original MS2×24 tag and allows single molecule visualization for extended periods of times even at high frame rates^13^. Moreover, this construct bears the lower affinity variant of the MS2 stem-loops (U instead of C in third position of the loop), providing excellent RNA visualization while preserving normal RNA degradation^24^.

**Figure 2:**
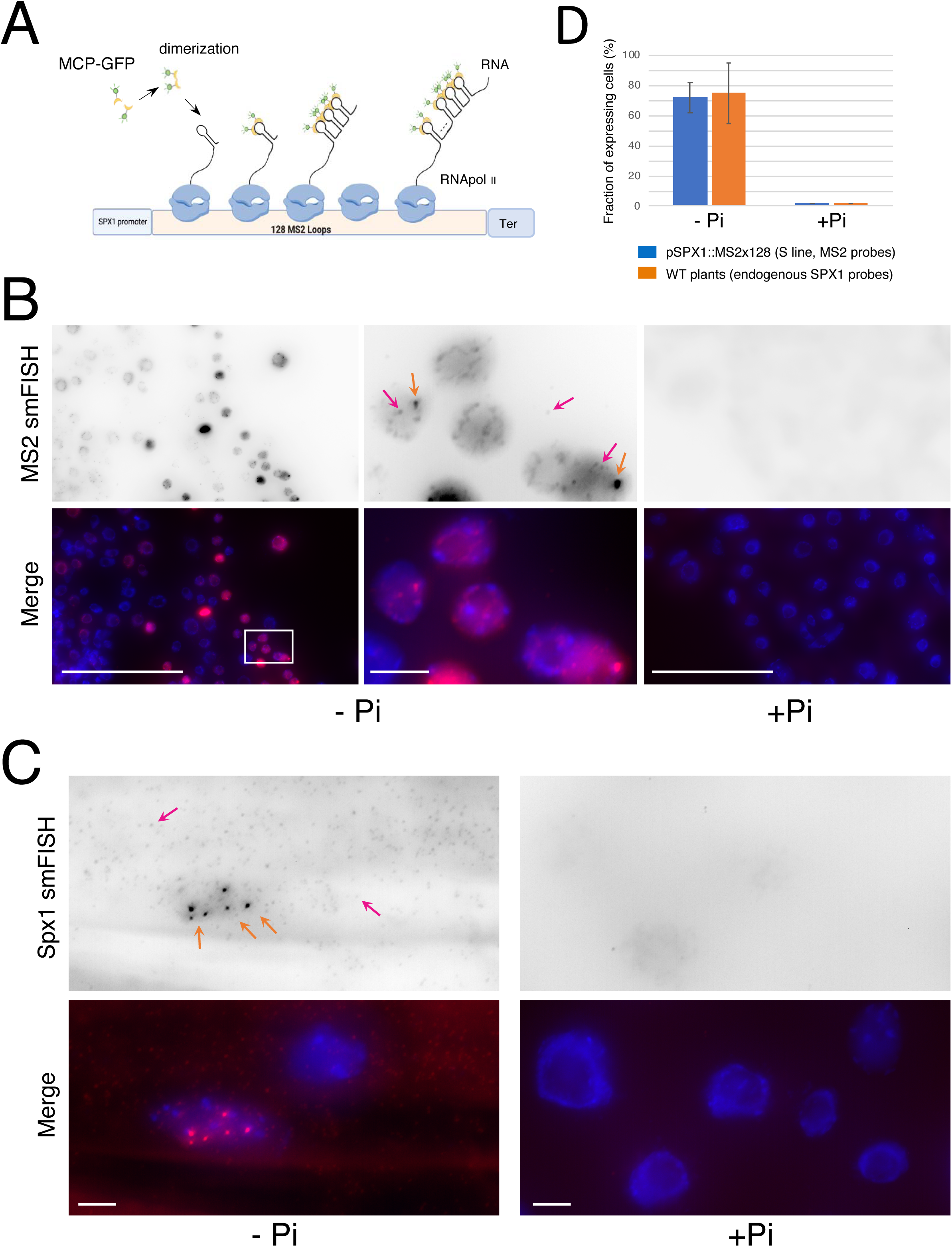
Validation of pSPX1::MS2×128 transgenic plants. (A) Principle of the MS2-MCP system. The transgene is under the control of the *SPX1* promoter and synthesizes a reporter RNA bearing 128 MS2 stem-loops, which are recognized by the MCP protein fused to a fluorescent marker (eGFP). (B) Microscopy images of squashed root caps of the transgenic S line expressing pSPX1::MS2×128 and processed for smFISH with probes hybridizing to the MS2×128 sequence. Left: Pi depleted sample (-Pi) and zoomed over the boxed area in the middle panels. Right: sample grown on Pi rich medium (+Pi). Top: smFISH signals; bottom: smFISH signals colored in red and merged with Dapi (blue). Images are maximal projections of z-stacks (widefield microscopy). Scale bars: 40 µm (left and right panels), and 4 µm (middle panels). Transcription sites and single RNA molecules are indicated by orange and pink arrows, respectively. (C) Legend as in B except that smFISH was performed with probes hybridizing against the endogenous *SPX1* mRNAs in WT squashed roots in the cap (right panel) and mature tissue (left panel). Scale bars: 4 µm. (D) Bar plot depicting the number of cells expressing the endogenous *SPX1* mRNA (orange, WT plants), or the pSPX1::MS2×128 reporter RNA (blue, S plants) in root (data for root cap and mature root area are combined), with (+Pi) or without (-Pi) phosphate. Mean and standard deviation estimated from 12 fields of view for *SPX1* probes (n=300) and 23 for MS2 probes (n=3087). Expressing cells are defined here as cells having at least 20 RNA molecules in the nucleus or the cytoplasm.

We developped Moclo vectors adapted to the Golden Gate system^25^ to facilitate the cloning and use of the MS2×128 recombinogenic sequence (Fig S2A), and we generated transcriptional fusions with the *SPX1* and *UNI1* promoters (Fig. 2A). We then introduced into the binary vector used for plant transformation a gene expressing a nuclear targeted MCP-eGFP fusion protein^13^ under the control of the weak constitutive Ubiquitin-10 promoter^26^ (Fig S2A). For the two constructs, genetic analysis selected homozygous transformants exhibiting single locus insertions. We first focused on the homozygous pSPX1::MS2×128 line named ‘S’ for strong and exhibiting the best signals. Molecular analysis revealed the insertion of three copies of the transgene at a single locus in the S line (Fig S3). The presence of the cap and the polyA tail, crucial elements for RNA stability, was also verified for MS2×128 transcripts (Fig S4). Single-molecule fluorescence in situ hybridization (smFISH) was performed on root squashes to analyze the phosphate response of the transgenes. Root squashing allows good probe penetration in tissues and causes some cells to fall off the root and adhere to coverslips as a monolayer^27^, allowing high quality imaging using wide-field microscopy with high numerical aperture objectives (Fig. 2B and S2B). The fluorescent oligonucleotide probes hybridizing against the MS2 stem-loops labelled the reporter RNA in most tissues of the root grown on Pi depleted medium, including the root cap and mature tissues (Fig. 2B and see below Fig. 3), and they did not detect signals in Col0 negative controls lacking the transgene (Fig. S5). Interestingly, the smFISH signals were rarely detected in the cytosol and mostly stained nuclei where transcription sites were visible as bright spots. This nuclear localization could be due to either a nuclear retention of the MS2-labelled RNA, or a reduced stability in the cytosol. In any case, the smFISH signal disappeared when plants were grown in the presence of phosphate (Fig. 2B, right panels). Quantification of the signals indicated that 74% of the cells expressed the pSPX1::MS2×128 transgene in absence of phosphate (>20 RNA/cell), and none in its presence (Fig. 2B and 2D). To further validate these results, we visualized the endogenous *SPX1* transcripts with a mix of 24 fluorescent oligonucleotide probes specific for SPX1. In Pi depleted conditions, the probes identified the transcription sites and single mRNA molecules present in the cytoplasm, as expected for a translated transcript (Fig 2C and S2B). Similar to the pSPX1::MS2×128 reporter, the endogenous *SPX1* mRNA was expressed throughout the root, including the cap and mature tissues^28^, with a number of positive cells similar to that of the reporter (Fig. 2D). Importantly, *SPX1* smFISH signals were very low in plants growing on Pi rich medium (Fig 2D and 2C, right panels) and completely absent in a *spx1spx2* double mutant used as negative control (Fig S6). Overall, these data indicated that the MS2 transgene driven by the *SPX1* promoter faithfully reported on the expression of *SPX1*, with a similar regulation and spatial expression pattern.

**Figure 3:**
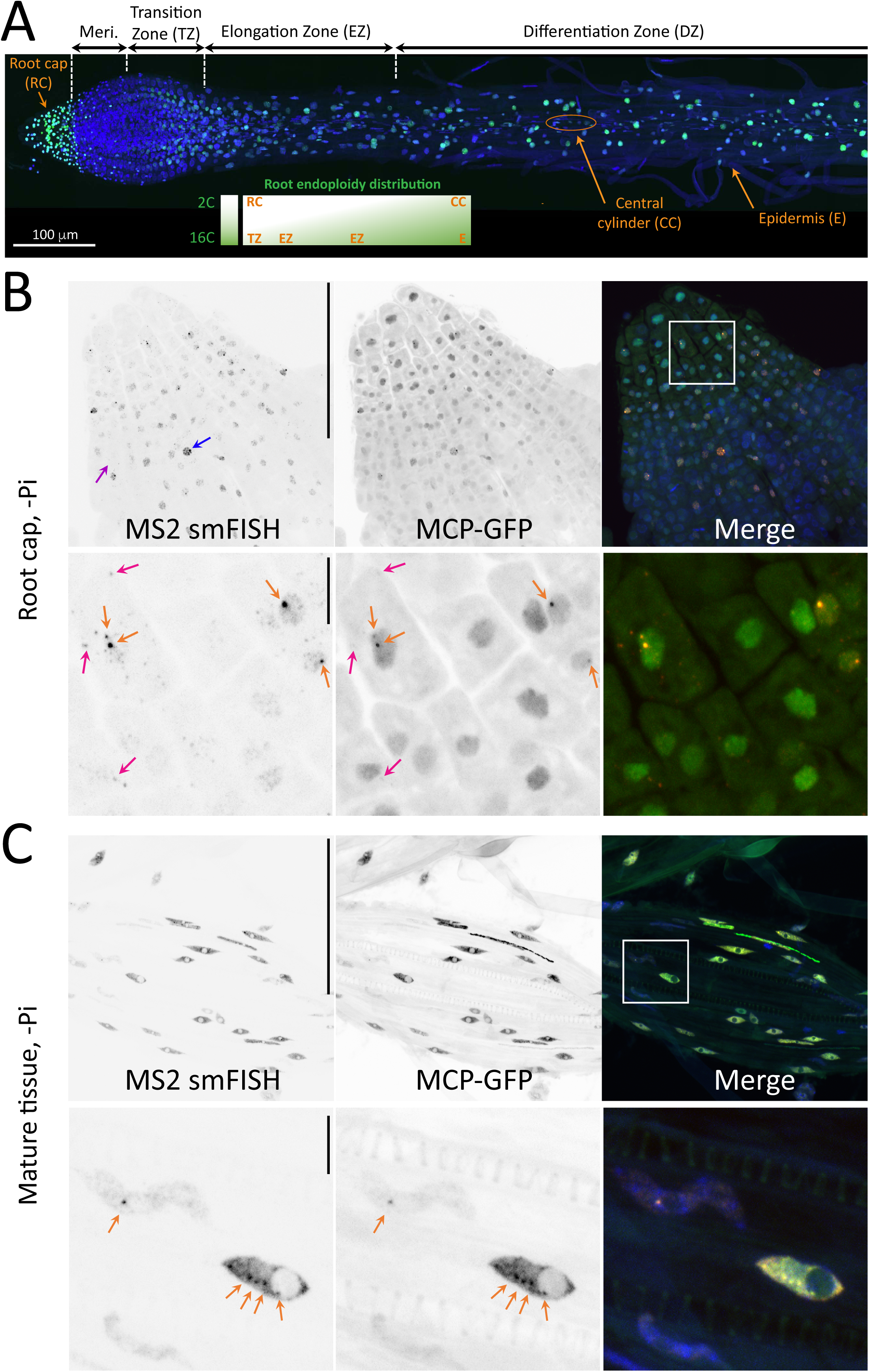
Imaging *SPX1* transcription in fixed plant tissues reveals allelic differences in the root cap and polyploid expression in mature tissue. (A) Organization of Arabidopsis root, with the name of the various root parts and tissues overlayed on a microscopy mosaic image of a homozygous S plant with MCP-GFP (green) and DNA (blue). Inset: ploidy of the root tissue. Note that a viewer with a zoomable high resolution version of the mosaic can be accessed at https://imjoy.io/lite?plugin=muellerflorian/hani-ms2:hani-ms2-sample-3. (B) Images are maximal image projections of z-stack from a root cap of a homozygous S plant grown without Pi and imaged by spinning disk microscopy in three colors. Left (and red in the Merge panel): smFISH signals obtained with probes against the MS2 repeats; middle (and green in the Merge panel): MCP-GFP signals. Blue: nuclei stained with Dapi. The bottom panels are zooms of the boxed area in Merge panel. Pink arrows: single RNA molecule; orange arrows: transcription sites. Scale bar: 100 µm (top panels) and 10 µm (bottom panels). (C) Legend as in B, except that a mature part of the root is imaged. Note that images in A, B and C come from different plantlets.

### Imaging transcription in fixed tissues of whole plants with MCP-GFP

We then turned to spinning disk confocal microscopy to image plants through the tissue depth (reaching 100µm) and to assess in detail the performance of MCP-GFP to image transcription. We first used the brightest homozygous pSPX1::MS2×128 line (called S) to image expression of the reporter in the root (Fig. 3A; a viewer with a zoomable high resolution image can be accessed at https://imjoy.io/lite?plugin=muellerflorian/hani-ms2:hani-ms2-sample-3 and https://imjoy.io/lite?plugin=muellerflorian/hani-ms2:hani-ms2-sample-1, which is a continuously growing organ with a well-defined architecture. At the apex, the root cap encapsulates the meristematic zone where cells divide rapidly until they enter the transition zone (TZ). Cells then elongate (EZ) and initiate tissue differentiation according to a well defined radial pattern, generating a number of different cell types (Fig. 3A).

First, we tested whether MCP-GFP could faithfully report on the MS2-tagged RNA and we performed smFISH together with MCP-GFP imaging. In the root cap, which is mostly composed of diploid cells^29^, smFISH against the MS2 stemloops revealed both single molecule and transcription sites, visible as bright spots in the nucleus (Fig. 3B, bottom panels; pink and orange arrows, respectively). In both root cap (Fig. 3B) and mature tissue (Fig. 3C), we observed colocalization between the smFISH signals revealed by MS2 probes and the fluorescent signal produced by MCP-GFP. The transcription sites were easily detected (Fig. 3B and C, orange arrows), and MCP-GFP could also reveal the brightest single mRNAs (Fig. 3B, pink arrows), indicating that the sensitivity of the system approches that of single molecules (see also below live cell experiments). To assess the reliability of detection, we counted transcription sites in both colors, and found that 90-95% of the sites detected by MCP-GFP were also labelled by MS2 smFISH.

Finally, we extended these observations to other *Arabidopsis* lines, using plants having either the pUnicorn1::MS2×128 reporter or another single locus transformant of the pSPX1::MS2×128 transgene (J line). The J line turned out to also present multiple T-DNA copies (Fig S3B), but it exhibited reduced levels of fluorescence when compared to the S line. The transcription sites of the reporters were readily detected with MCP-GFP and co-localized with the MS2 smFISH signals (Fig. S7). Overall, these results validated the MS2/MCP system as a robust and sensitive system to image transcription in whole plants tissues.

### Single molecule counting by smFISH gives insights into the dynamics of transcription in Arabidopsis *thaliana*

The ability of smFISH to detect single RNAs enables a quantitative analysis of the RNA polymerase II transcription cycle^9,12,30,31^. We therefore measured the brightness of transcription sites and used the visible single molecules to compute the absolute number of nascent RNA molecules at active transcription sites (Fig. 4A and B; see Methods). We obtained an average of 37 molecules at the transcription site of the pSPX1::MS2×128 reporter for the S lines, with a main peak at ∼20 molecules (Fig. 4A). We also designed a set of 21 fluorescent oligonucleotides against the sequence immediately downstream the polyadenylation site of the reporter (see Fig. S12). We however failed to detect any smFISH signal, indicating that 3’-end processing was rapid as compared to transcription. This agrees with previous GRO-seq experiments that revealed that RNA polymerases terminate transcription very rapidly after polyA sites^32,33^. We repeated these measurements for the endogenous *SPX1* mRNA and found an average of 11.3 molecules at transcription sites, with a main peak at 9 molecules. Given that the *SPX1* gene is 1.4 kb long, this number suggests that active *SPX1* genes have one polymerase every 77 bases if 3’-end processing is immediate, and one polymerase every 120 bases if 3’-end processing takes a minute^13^ (assuming an elongation rate of 2 kb/min^30^). For the 3kb pSPX1::MS2×128 reporter, similar numbers were found if one considers that the peak of 25 molecules at a transcription site corresponds to a single active copy (Fig. 4A). These numbers indicate that active *SPX1* genes have an initiation event every 2.3 to 3.5 seconds on average (see Methods and schematic in Fig. 4B). This is in the high range of previous estimates obtained in human cells and Drosophila^13^. It suggests that transcription in *Arabidopsis* is rapid and occurs in the form of polymerase convoys, produced by initiation events rapidly occurring one after another when the gene turns on^13^.

**Figure 4:**
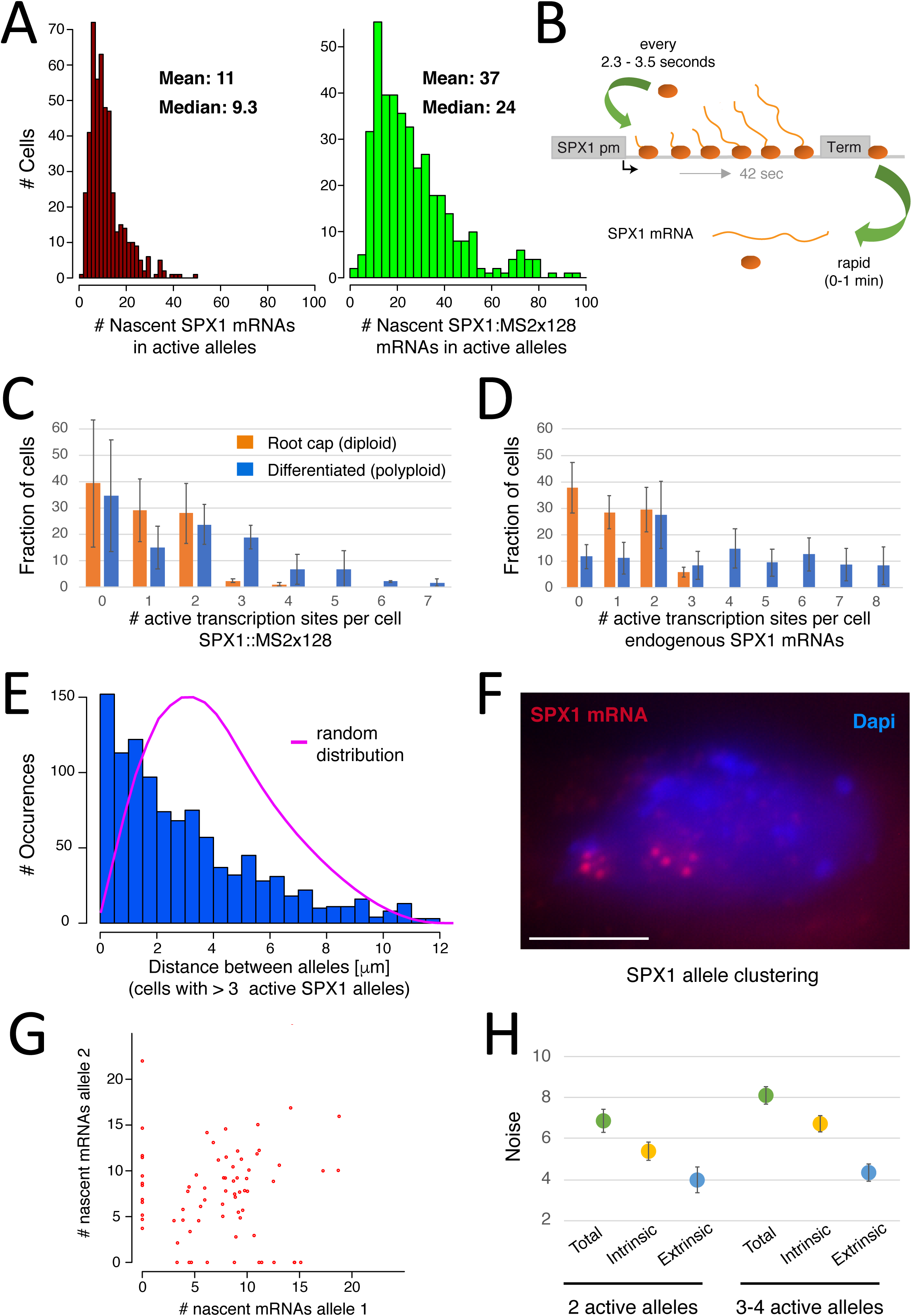
Quantitation of *SPX1* transcription in fixed cells gives insights into transcription dynamics, ploidy and intrinsic *vs.* extrinsic noise. (A) Graph depicting the distribution of the brightness of active transcription sites for the endogenous *SPX1* gene (left), or the SPX1-MS2×128 reporter (right) both detected by smFISH. Brightness values are expressed in number of full-length RNA molecules (x-axis), and the y-axis represent the number of cells with these values. (B) Model of *SPX1* transcription, based on smFISH data labelling the endogenous *SPX1* mRNAs. (C) Graph depicting the number of active transcription sites per cell for the SPX1-MS2×128 reporter detected by smFISH on 4 plants (488 nuclei) for mature tissue and 3 plants (641 nuclei) for root cap. x-axis: number of active transcription sites per cell; y-axis: fraction of cells with these values. Orange bars: diploid root cap cells; blue bars: differentiated, polyploid cells. Error bars represent standard deviation. (D) Graph depicting the number of active transcription sites per cell for the endogenous *SPX1* mRNA detected by smFISH on 9 plants (300 nuclei) for mature and 3 plants (137 nuclei) for root cap. Legend as in C. (E) Histogram depicting the distance distribution between active transcription sites in single cells, for the endogenous *SPX1* gene and for polyploid cells having 4 or more sites. Distances are expressed in 1m (x-axis), and the y-axis represent the number of cells with these values. Pink curve: distance distribution obtained by simulating a random location of 8 spots within Arabidopsis nuclei. (F) Image is a maximal image projection of a z-stack from a differentiated root cell labelled by smFISH with probes against the endogenous *SPX1* mRNA (widefield microcopy). Red: smFISH signals; blue: nucleus stained with Dapi. Scale bar: 5 µm. (G) Graph depicting the correlation of activities of *SPX1* alleles, in cells having one or two active transcription sites. x-axis: number of molecules at the first transcription site; y-axis: number of molecules at the second transcription site. (H) Graph depicting the levels of total, intrinsic and extrinsic noise for cells with either 2 active transcription sites (left), or 3-4 (right). Error bars represent the error estimated by bootstrapping (see Methods).

### Quantitation of *SPX1* transcription in fixed roots reveals cell polyploidy in mature tissue as well as large allelic differences

Endoreplication occurs frequently in root with DNA cell contents ranging from 2C to 16C^29^. This phenomenon increased as one moves away from the root tip. Being involved in the size of the cells, it also varies and increases from the central to the expended external cell layers such as cortex or epidermis^29^ (Fig. 3A). The number of transcription sites present in each nucleus highlighted this endoreplication. In the root apex of the pSPX1::MS2×128 S line, cells of the columella (extremity of the root cap) exhibited in the vast majority of cases no more than two transcription sites for the MS2 reporter or the endogenous *SPX1* gene, as expected for a mainly diploid tissue^29^ (Fig. 3B and Fig. 4C for quantifications). In contrast, images recorded in the differenciated part of the root revealed the more complex nature of older tissues with the presence of cells with many transcription sites, indicating polyploidy (Fig. 3C and 2C). Quantification of the number of active alleles per cell indicated that in diploid columella cells, only 3% of the cells had more than 2 transcription sites, while in the mature tissue, 35% had more than two sites, with 2% having as many as 7 (Fig. 4C and 4D). Similar numbers were obtained with the endogenous *SPX1* gene (Fig. 4D). Interestingly, we noted that in cells with many active alleles, the *SPX1* transcription sites frequently clustered in the nucleoplasm, most often forming two groups (see Fig. 4F for an example). To explore this further, we measured the distance between all visible transcription sites in cells having 4 to 8 active sites, and we compared the resulting distance distribution to a situation were we simulated 8 sites with a random location (Fig. 4E, see Methods). This showed that the transcription sites had a non-random distribution and were frequently close to one another, with 40% of the distances falling within 1.5 microns. This phenomenon was also visible when we compared the brightness of transcription sites in cells having one or two transcription sites, to cells having four or more sites (Fig. S2C). Cells with one or two SPX1 transcription sites had a single peak of 10 nascent RNAs, while cells with four or more sites had a second peak at 20 nascent RNAs (Fig. S2C). This suggests that in these cells, some transcription sites had coalesced into a single spot, reminiscent of the ‘transcription factories’ previously described in human cells^34^.

Next, we compared the activities of the different alleles of single cells, focusing first diploid root cap cells. Surprisingly, an important number of cells exhibited only a single active transcription site (30 % of the cells; Fig. 4C), or no transcription at all (39%). This was also observed with the endogenous *SPX1* gene (Fig. 4D), indicating that it is a feature of this Arabidopsis gene and not an artifact of the MS2 reporter. Since the cells with no transcription sites contained smFISH signal in the nucleoplasm or the cytoplasm (Fig. 3), transcription had been active in these cells, therefore indicating that *SPX1* promoter activity was discontinuous. Such discontinuous transcription is the result of gene bursting and has been oberved in many organisms including yeast, Drosophila and mammals^35^. To our knowledge, it has not been reported so far in plants. Gene bursting involves the stochastic switching of a promoter between active and inactive states. It depends on mechanistic aspects of transcription initiation as well as transcriptional regulation^12^. This stochasticity causes variations in gene expression among identical cells (ie. gene expression noise), which has sometimes important phenotypic consequences^12^. Commonly, two sources of noise are distinguished: intrinsic and extrinsic. Intrinsic noise is due to the stochastic nature of biochemical reactions involving a single molecule of DNA and the transcription factors acting on it, and it occurs independently on each allele. In contrast, extrinsic noise modulates similarly both alleles as the result of events affecting the entire cell (such as cell cycle or activation of a signaling pathway). Interestingly, our capability of accessing several alleles within a cell raised the possibility to discriminate between the two causes of transcriptional noise. For each cell that had only one or two active alleles, we plotted the brightness of one *SPX1* allele as a function of the other (Fig. 4G). This revealed both correlated (cells on the diagonal) and uncorrelated (cells off the diagonal) transcriptional activities of these alleles. To quantify this phenomenon further, we measured the total, intrinsic and extrinsic noise^36,37^, using cells having either exactly two, or 3-4 active transcription sites (Fig. 4H, see Methods). As expected from the results described above, we found that the total noise had significant intrinsic and extrinsic contributions, with intrinsic noise being the dominant source. This highlights the quantitative importance of transcriptional noise for Arabidopsis.

### Real-time visualization of SPX1 transcription reveals gene bursting and Pi mediated repression

We used a simplified version^38^ of the RootChip microfluidic system^22^ to combine live-cell imaging with the capacity to change the phosphate solution rapidly (Fig. 5A and Movies). Imaging was performed with spinning disk microscopy, which offered the best compromise between image quality, size of the field of view and image acquisition speed. A typical experiment recorded at each time point a z-stack of 200 images with a z-spacing of 500 nm, allowing us to image at high resolution the entire root and thus access all its tissues. One image stack was recorded every 2-3 minutes for ∼1 hour, and maximum intensity projection produced clear images where multiple transcription sites could be analyzed. Using the brightest S line expressing pSPX1::MS2×128 and grown without Pi, images from the first time point confirmed the transcriptional heterogeneity between cells, with neighboring cells exhibiting zero, one, two or even more active transcription sites and with the intensity of transcription sites varying by a factor of 4 (Fig. 5-7 and Movies 3-7). Observation of the root constantly supplied with -Pi solution directly demonstrated bursting of the SPX1 promoter, with transcription sites coming on and off over periods of minutes (Fig. 6, Movies 1 and 2). Interestingly, this phenomenon was mainly observed in the root cap, and only very rarely in mature tissues. Remarkably, the arrest of transcription after a burst provided an opportunity to visualize the release of single RNAs from the transcription sites (Fig. 6D, Movie 2), highlighting the dynamics of the process and confirming single molecule sensitivity in live plants.

**Figure 5:**
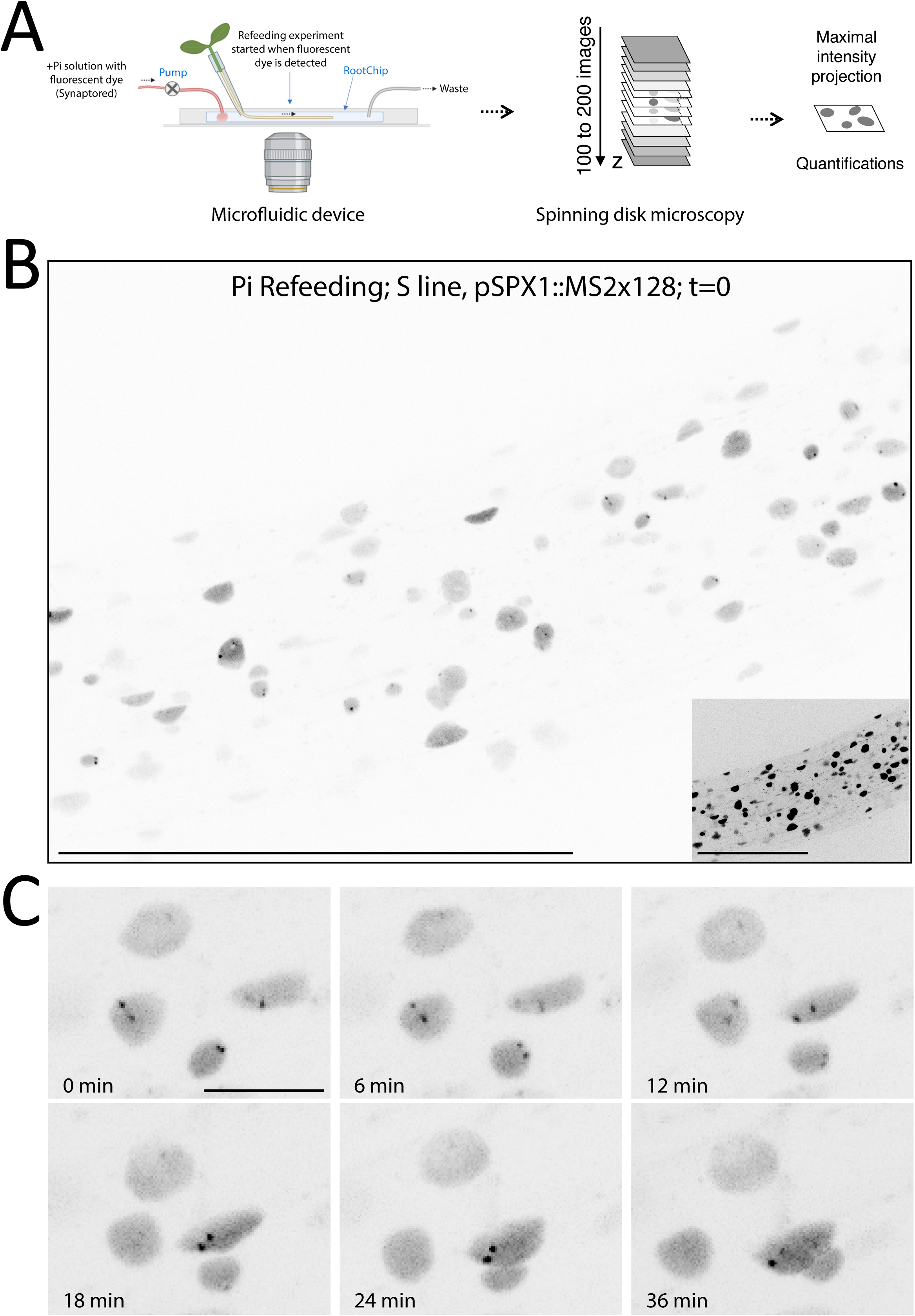
Combining microfluidic and MS2 technology reveals the fast transcriptional repression triggered by Pi supply in pSPX1::MS2×128 transgenic plants. (A) Principle of experiment combining RootChip microfluidic system with spinning disk microscopy to analyze the transcriptional response to Pi refeeding in real time. (B) Image is a maximal image projection from a time-lapse movie recorded in 3D (200 z-planes), which displays MCP-GFP fluorescence in S plants at the start of Pi refeeding (t=0). Scale bar: 100 µm. Inset: identical image with a higher contrast to display the tissue structure. (C) Images are maximal image projection from the time-lapse movie shown in B, taken at the indicated time points after Pi refeeding and displaying MCP-GFP fluorescence. Scale bar: 10 µm.

**Figure 6:**
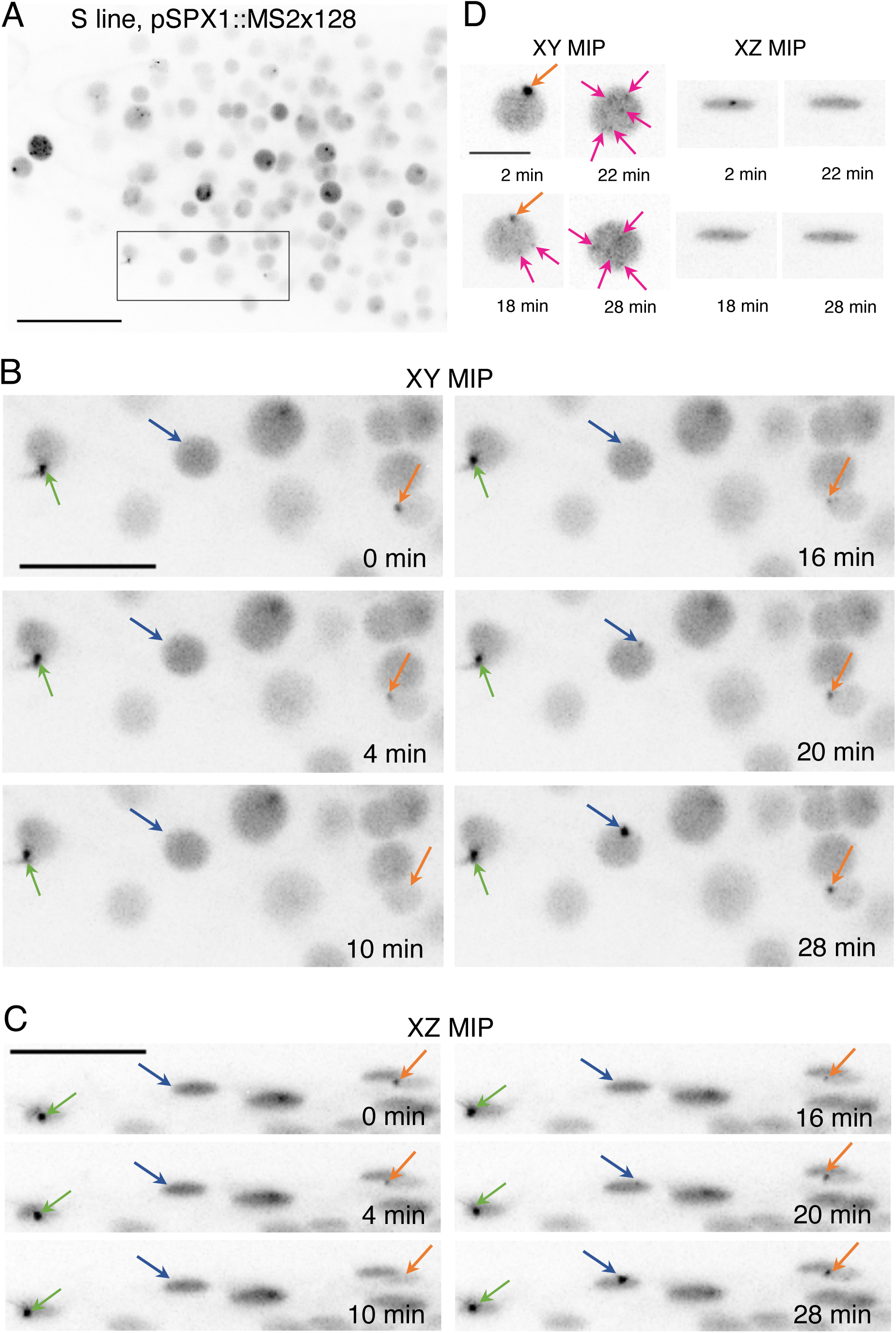
The SPX1 promoter generates bursts of activity in root cap cells grown at steady-state in absence of phosphate. (A) Image is a maximal image projection from a time-lapse movie recorded in 3D (44 z-planes 600 nm apart), which displays MCP-GFP fluorescence in S plants. Scale bar: 20 µm. (B and C) Images are maximal image projection (MIP) on XY (A) or XZ (B) axes from the time-lapse movie shown in A, taken at the indicated time points and displaying MCP-GFP fluorescence. Arrows pinpoint transcription sites where the signal remains constant (green), increases (blue) or turns off and then on (orange) during time-laps. Scale bar: 10 µm. (D) Images are maximal image projection (MIP) on XY (A) or XZ (B) axes of a nucleus where transcription decreases at transcription site (orange arrow) and individual mRNA released can be seen (pink arrows). Scale bar: 5 µm.

**Figure 7:**
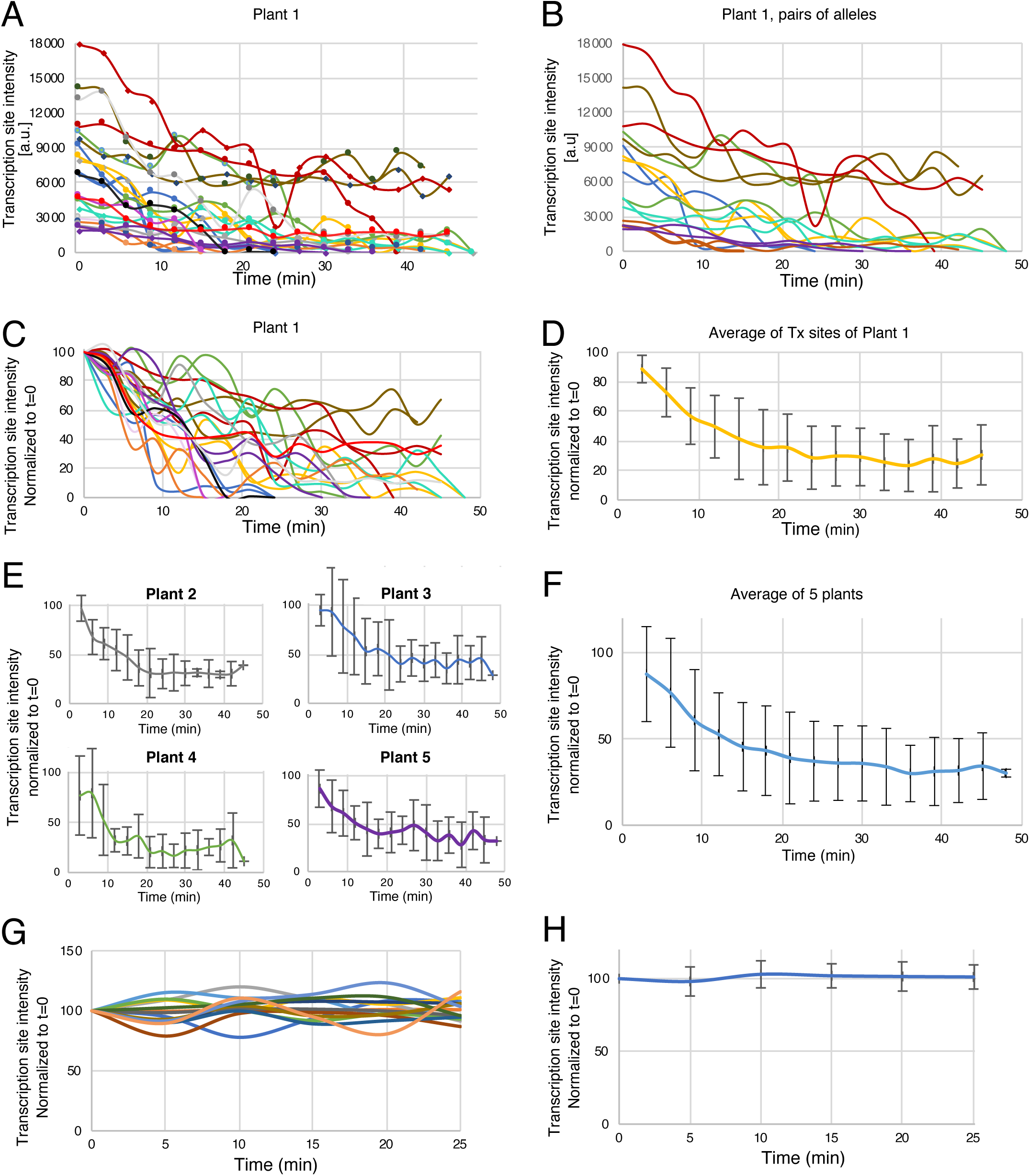
Analysis of transcription site activity following Pi supply in pSPX1::MS2×128 S line transgenic plants. (A) Intensity of fluorescence signals at transcription sites in arbitrary units (a.u.), and recorded every 3 minutes after Pi refeeding, for cells exhibiting one or two active transcription sites (n=22) recorded in 14 mature root cells located 1mm above the root tip. (B) Intensity of fluorescence of transcription sites over time, for 8 cells exhibiting two active transcription sites (plotted in same color). (C) Normalized intensities of transcription sites over time, for the same cells as in A. Normalization is done according the value measured at t=0. (D) Average normalized intensities of transcription site, for the 22 transcription sites shown in panels A and B (mean and standard deviation). (E) Same average curve of normalized transcription site intensities as (D) obtained with independent plant samples. (F) Average curve obtained by analysis of 67 transcription sites analyzed on 5 plants. (G) Intensity of fluorescence signals at transcription sites in arbitrary units (a.u.), and recorded every 3 minutes for cells exhibiting one or two active transcription sites of plant supplied with Pi depleted nutrient solution, (n=14) (H) Normalized intensities of transcription sites over time, for the same cells as in G. Normalization is done according the value measured at t=0. Standard deviation is provided (n=14).

Next, we analyzed the response to Pi resupply. To this end, the S line was first grown during a few days in -Pi liquid medium, before receiving phosphate rich medium combined with the Synaptored^TM^ fluorescent dye. This allowed to precisely record the arrival of the supplemented media on the imaging field, thereby defining the time 0 of the time course. The movement of nuclei in the cells and the displacement of cells themselves complicated transcription site tracking. To simplify the analysis of the time course, we therefore used only cells with one or two active transcription sites. The repression of transcription upon Pi resupply proved to be rapid (Fig. 5 A-C, S8, S9 and Movies 3-7). Signals were then normalized to the value obtained at time 0 (Fig. 7C) and an average response curve was produced (Fig. 7D). The measurements were repeated with independent samples and they all showed similar curves with a signal starting to decrease between 0 to 6 minutes after Pi provision at the root surface (Fig. 7E). Similar results were obtained with independent transgenic J line (Fig. S10, Movie 8), revealing the extraordinary sensitivity and rapidity of the regulatory cascade triggered by Pi.

The decrease of fluorescence starting with Pi resupply reached the root continued until it reached a plateau 18 to 25 minutes later (Fig. 7F and S10). Interestingly, cells presenting multiple transcription sites revealed on average a close coordination of the repression between these sites, indicating that extrinsic factors dominate gene regulation when Pi repression initiates, as expected for a signaling pathway affecting the entire cell (Fig. 5C, 7B and S8). Controls continuously supplied with Pi depleted solution in mature tissue did not show significant changes (Fig. 7 G, H and S11) over 25 min observation period.

The comparison between the MS2 and RTqPCR results revealed a delay of 20-30 minutes for the RT-qPCR to show the maximum repression, which is in agreement with the *SPX1* mRNA half-life previously estimated in the range of 15-20 min (Fig. 1A and S1A,B). Overall, these results demonstrate the unexpectedly fast dynamics of the plant response to Pi resupply, and nicely illustrates the capacity of the MS2 system to access transcriptional regulation in live plants. It should be noticed that in mature tissues, we restricted the analysis to nuclei exhibiting one or two active alleles to avoid misattribution during the analysis (resulting from nuclei movement during analysis). This favors more central cell (less affected by polyploidy) representation, which are known to be less affected by Pi repression compared to cortex or epidermis cells^39^. This may explain the slight difference (around 10%) of average extent of the decrease in expression between RTqPCR and MS2 live imaging.

## Discussion

### Large MS2 arrays and microfluidics reveal single cell transcriptional dynamics in live plants

Fluorescent RNA technologies provide access to single molecule studies offering invaluable insights into gene expression mechanisms^12,40^. A previous attempt to use these technologies in plants studied a highly abundant viral RNA in transient transformants and low signal-to-noise ratio prevented their use in stable plant lines^41^. The MS2×128 construct developed here (which uses six time more MS2 loops) solves this problem and provides a direct acces to transcriptional activities of live plants, at the level of single alleles and with a sensitivity nearing single RNAs. Nevertheless, single molecule detection remains difficult and not exhaustive, due in particular to the necessity of limiting illumination power to prevent bleaching and phototoxicity when acquiring 200 z planes for tens of time points. We hope that new constructions currently underway, where the number of MS2 loops is doubled (256) or MCP-GFP expression is optimized, will provide important future improvements. Originally developped for yeast or mammalian cell lines, fluorescent tagging of RNA has been introduced in only few multicellular organisms (Drosophila embryos in particular^10,35^), mostly to study transcriptional regulation during embryonic development and in absence of external perturbations. Here, the implementation of microfluidics allowed a complete control of environmental conditions and the investigation of the root response to phosphate starvation and resupply. Measurements of transcriptional inhibition often suffer from both indirect measurements introducing a considerable lag time in repression detection, and ensemble measurments averaging effects. In contrast, the tools developed here allow visualization of repression in real time and single cells, and demonstrate that it is a very rapid process that takes place in minutes.

Plants are sessile organisms that developed exquisitely efficient regulation systems to ensure homeostasis in face of changing environmental conditions. Due to the fast and sensitive tuning of transcriptional mechanisms, microfluidics offers the ideal solution to control environmental conditions. This strategy could be extended to a variety of other stresses, including biotic ones, and to all plant organs as various microfluidic systems exist to study aerial parts. The technological developments made here thus open entirely new possibilities by allowing the direct visualization of transcriptional regulations in living plants, in real time and at the level of single cells and single molecules.

### Integration of single cell responses to phosphate signaling at the tissular level

The capacity to dissect transcriptional responses of individual cells within a tissue and between different tissues within an organ is a major step forward. The use of real-time phosphate radioisotope imaging revealed that Pi enters and accumulates in the root tip within less than one minute ^42^. The present study thus directly links this arrival with transcriptional repression, demonstrating the close concomitance between the two phenomena. Moreover, our analysis, owing to unprecedented cellular resolution in plants, also identified the heterogeneity of transcriptional responses. Indeed, we observe strikingly different responses to phosphate resupply in different tissues of the root: a cell can repress *SPX1* transcription within minutes while its neighbor may continue transcribing unabatedly (Fig. 3B, 5C and 7). The fact that the alleles of a cell show a coordinated response indicates that this heterogeneity in repression arises mainly from extrinsic sources, as expected for a regulation involving a cellular signaling pathway. Interestingly, the cell-to-cell heterogeneity of *SPX1* transcription is already visible at steady state when plants are starved from phosphate: a third of root cap cells contain *SPX1* mRNAs while not being transcriptionally active, indicating discontinuous promoter activity. This peculiar phenomenon of gene bursting is an important driver of cell heterogeneity and has already been described in animals^35^, but not plants. Our analysis suggested that this phenomenon may differ greatly between cell types. Using specific cell layer markers in the future will help to investigate more precisely this phenomenon and decipher putative driving mechanisms. We show here with *SPX1* that transcriptional noise in plants has both intrinsic and extrinsic components, suggesting that it arises from both stochastic promoter dynamics and cellular regulatory pathways. At steady state, in absence of Pi, intrinsic noise appears to be dominant, while extrinsic factors appear to take over during Pi refeeding, generating heterogeneity in repression kinetics and in the extent of repression. This needs to be investigated in different organs, developmental stages, physiological statuses and environmental contexts. Because of its role in cell heterogeneity, transcriptional noise is clearly a key point to understand how the activity of individual cells is integrated within entire tissues.

### Transcription imaging in polyploid cells reveals clustering of active alleles

Endoreplication in plants has been associated with high metabolic activity or fast biomass development, economically important for most vegetables and fruits. Recent analysis of Arabidopsis illustrated the extent of this process affecting many root cells during their development and differentiation, the highest level of endoploidy (16C) being observed either in most developped cells (epidermis hair cell) or in highly active regions associated with distribution of nutrients (phloem companion cells)^29^. So far endoploidy detection uses destructive techniques such as flow cytometry or histochemical staining of endocycles markers or DNA. Here, we overcame such limitations providing the opportunities to access kinetics of endoploidy, taking into account development, cell fate, biotic or abiotic stresses. Our results highlighted the magnitude of somatic polyploidy in a root, revealing that many alleles of polyploid cells can be simultaneously transcriptionally active. We found that in absence of phosphate, active *SPX1* alleles of polyploid cells tended to cluster together, possibly because endoreplicated chromatids generally occupy the same chromosome territory^43^. More surprisingly,our data also suggest that active SPX1 alleles have a propensy to coalesce together, a phenomenon reminiscent of transcriptional factories reported in animal cells^34^. In the future, it will be important to confirm the coalescence of active alleles by directly observing this phenomenon in live cell. It will be also interesting to determine whether this is related to the very high transcriptional activity of the *SPX1* gene in these conditions (one initiation event every 2-4 seconds), and to understand how this process occurs and whether it contributes to gene regulation. Note that these initiation rates are based on polymerase elongation speed measured in animals or yeast. They do not take into account possible modifications affecting transcription elongation, pausing and termination, which are poorly described in plants despite recent advances^32,33^, and will require additional experiments to account for them.

Overall, the technology presented here offers direct access to the RNA polymerase II activity in live cells and could be used to study crucial plant phenomena affecting transcription (silencing, heterozygosity, impact of gametophyte origin …). This unprecedented spatio-temporal resolution for plant transcription therefore provides new horizons to multiple applications hitherto inaccessible for plant physiology.

## Material and Methods

### Plant materials and growth conditions

Wild type Arabidopsis thaliana Col-0 seeds were sterilized and grown vertically on Murashige and Skoog medium diluted 10-fold (MS/10) in Petri dishes supplemented with a Pi source containing either 500 µM (+P) or 13 µM (-P) KH2PO4 in a culture chamber under a 16-hr-light/8-hr-dark regime (25°C/22°C), as previously described^44^.

For global transcriptomic analysis, 7 days old seedlings were germinated on a sterile nylon mesh deposited at the surface of the culture medium to facilitate the transfer between media. For resupply experiment, plantlets were transferred from -P to +P for 30, 60 or 180 minutes. In order to minimize stress related transfer, control plants in -P and +P media were also transferred for similar period of time on same media.

In order to get closer to microfluidic conditions and limit the stresses related to the solid media transfer, we modified the protocol for subsequent experiments (RTqPCR analyses), by transferring the 7 days old seedling from agar plate to liquid MS/10 for 3 days. Then, -P plants were resupplied with +P solution for up to 3 hours before collecting the roots for RNA extraction. Transcription inhibition experiments were performed by addition of 0.6 mM cordycepin ^45^. For each condition three independent biological replicates were analyzed. Plant DNA extraction and genetic segregation analysis were performed as previously described^46^.

### RNA-seq library preparation and analysis (RNA seq and RTqPCR)

The extraction of total RNA from Arabidopsis roots or shoots, the synthesis of RNA-seq libraries and their analysis using Illumina sequencing technology (Illumina, San Diego, CA) were performed as previously described ^47^. Quantitative RTqPCR experiments were performed as previously described ^48^ with primers listed in the Supplemental Table 1.

For 7-Methylgunaosine RNA Immunoprecipitation (m7G-RIP) experiment, total RNA was extracted using Monarch Total RNA Miniprep kit including DNAse treatment on column (New England Biolabs). m7G-RIP was then performed as previously described^49^ with slight modifications. 5 μg of total RNA was used as starting material. Elution was performed using Guanidium 8M 10 minutes at 65°C followed by RNA precipitation. As negative control, pyrophosphatase treatment was performed on total RNA prior to m7G-RIP. Reverse-transcription was then performed on input and eluate fractions using 500 ng of RNA and oligo dT primer. PCR was then performed using primers against MS2 transgene or endogenous SPX1 using oligos described in Supplemental Table 1.

### Transgene constructs

The pSPX1::MS2×128 and pUNI1::MS2×128 constructs were built using the Golden gate cloning technique ^25^. MS2×128 stem loops (pMK123-MS2×128-XbaI) and NOS terminator were inserted into level 0 vectors pICH41308, and pICH41421 respectively. The promoters of *SPX1* (At5G20150) and *UNICORN1* (At5g20790) were PCR amplified (1452 bp^28^ and 2161 bp upstream of the start codon) using oligos described in Supplemental Table 1. These sequences were introduced in level0 vectors (pICH41295). The levels 0 modules were assembled directly into a level 2 expression vector pICH86966.

The Ubiquitin10 promoter (At4g05320) driving eGFP expression derived from pUBN-Dest^26^ was assembled with MCP contained in pDONR201^8^ using the Gateway cloning technique (Invitrogen) creating plasmid pUb::eGFP/MCP-NLS. This construct was excised from the Gateway vector using PmeI and SmaI (Invitrogen) and introduced in level2 T-DNA pSPX1::MS2×128 or pUNI1::MS2×128 vectors using PmeI restriction sites. A clone having the two constructs in opposite orientations creating pMCP/proSPX1::MS2 was used to transform Agrobacterium tumefaciens strain C58C1. Figure S2A summarizes the cloning strategy. The reporter line carrying Luciferase in fusion with the UNICORN1 promoter was generated as previously published ^44^.

### Production of transgenic lines

Transformants produced by floral dipping^50^ were selected on Hoagland/2 media complemented with 50 mg/L kanamycin. The progeny exhibiting 3:1 segregation were carried to T3 generation to identify homozygous lines carrying a single insertion locus. At least 5 independent lines were obtained for each construct.

### Single molecule Fluorescence *In Situ* Hybridization

*Arabidopsis* seedlings were grown on +P and -P agar media. The roots of 7 to 10 day old seedlings were collected and fixed for 30 min with 2-(*N*-morpholino) ethane sulfonic acid (MES) buffer solution pH 5.7 containing 4% paraformaldehyde (32% methanol-free solution; Electron microscopy Sciences). pH was adjusted to 5.7 with KOH solution.

Roots were rinsed twice with MES buffer (pH 5.8) and put onto a microscopic slide with a coverslip. They were smoothly squashed (to splay them and produce a single cell layer) and submerged briefly in liquid nitrogen. Then, after removing coverslips, they were left out to dry at room temperature for one hour. To permeabilize the samples, the slides were immersed in 70% ethanol overnight on a rotary shaker prior to hybridization. The ethanol was evaporated at room temperature before washing the roots with MES.

SmFISH was used for the detection of MS2 repeats and endogenous *SPX1* mRNAs. The MS2 probe was made of a mix of 10 pre-labelled fluorescent oligos directed against 32xMS2 repeats ^13^. Each oligo contains 2 to 4 molecules of Cy3 and hybridizes 4 times across 128xMS2 repeats, allowing the binding of 40 probes to each single RNA molecule. The set of *SPX1* probes was made of a mix of 24 fluorescent oligonucleotides carrying 2 to 3 Cy3 fluorophores and covering the entire SPX1 transcript including 5’ and 3’ UTR (Table S2). The pre-hybridization was performed in 1xSSC/40% formamide buffer for 15 min at room temperature. Hybridization was performed with 2 ng/µl probe and in 40% formamide for MS2, whereas for *SPX1* only 15% formamide was used. The smiFISH probe mix directed against the post-polyA region of the MS2 construct was made of 21 oligonucleotides (Table S2) and hybridized as previously described ^51^. After addition of probes, samples were covered with a coverslip and remained overnight at 37°C as previously described ^51^. For rinsing, after coverslip removal, root samples were washed twice during 45 min at 37°C with freshly prepared 1xSSC/40% formamide buffer and rinsed at room temperature with MES pH 5.8 buffer and dried. A drop of Prolong Diamond antifade mounting medium (Invitrogen) containing DAPI was added prior to observation. For long-term storage, slides were kept at -20°C.

### Fluorescence imaging of fixed plants

SmFISH and MCP-GFP images were taken using either a spinning disk confocal or a wide field microscope. For spinning disk microscopy, we used a Dragonfly (Oxford instrument) equipped with four laser lines (405, 488, 561, 637 nm) and an ultrasensitive EMCCD camera (iXon Life 888, Andor) mounted on a Nikon Eclipse Ti2 microscope body, using a 40x, NA 1.3 Plan Fluor oil objective or a 60x, NA 1.4 Plan Apochromat oil objective coupled with a supplementary lens of 2x, using z-stacks with a 0.5 1m or 0.4 1m step. For widefield imaging, we used a Zeiss Axioimager Z1 widefield upright microscope equipped with a camera sCMOS ZYLA 4.2 MP (Andor), using a 100x, NA 1.4 Plan Apochromat oil objective. For these z stacks, a step of 0.3 or 0.4 1m was used. Maximal image projections (MIP) were generated with ImageJ, and figures were realized using Adobe Photoshop and Illustrator. The mosaic of Figure 3A is accessible with a viewer run with Imjoy^52^. Note that plant fixation and smFISH both reduced the GFP signals.

### Analysis and quantifications of smFISH images

Quantification of the brightness of transcription sites was made with a modified version of HotSpot ^40^. This is a user-friendly MatLab software that allows to quantify smFISH signals in very noisy images, unlike FISH-quant^53,54^. Briefly, the user can navigate in 2D or 3D bicolor images to manually select transcription sites and single RNAs, and the script finds the local maxima and fits 2D or 3D gaussians to the selected transcription spots. The images of single RNAs are averaged and also fitted to 2D or 3D gaussians, and the integrated intensity of the transcription sites are divided by the intensity of average of single RNAs, yielding the exact number of molecules present at transcription sites. In this work, quantifications were done in 2D on MIPs using 100-500 single RNA per image to generate the average. HotSpot is available on GitHub (https://github.com/muellerflorian/hotspot).

To calculate initiation rates, we assumed that initiation occurs at random with a constant rate. This produces a uniform distribution of RNA polymerases along the *SPX1* gene following arithmetic series. Because the MS2 and *SPX1* probes are distributed homogeneously along the corresponding pre-mRNAs, incomplete nascent RNAs are labelled with only a fraction of the probes. If one assumes that the pre-mRNAs immediately leave the transcription site once polymerases reach the 3’-end of the gene (in agreement with experimental data), the barycenter of the polymerase distribution is the middle of the gene, and the average brightness of a nascent pre-mRNA is thus half that of a full-length pre-mRNA. In this case, the number of polymerase on the gene, Npol, is thus twice the brightness of transcription sites, expressed in number of full-length pre-mRNA (noted TS_RNA_). Assuming an elongation rate of 2kb/min, the time to transcribe the gene, t_el_, is 42 s, and the average initiation rate is Npol divided by t_el_. If the pre-mRNA takes t_proc_ seconds to be 3’-end processed and released from the transcription site, then Npol is TS_RNA_*[2*t_el_/(t_el_+t_proc_)+ t_proc_/(t_el_+t_proc_)], and the initiation rate is Npol divided by (t_el_+t_proc_).

To analyze the distribution of distances separating active transcription sites within single nuclei, we computed for each nuclei all the distances between all the visible transcription sites. To simulate a random distribution of transcription sites, Arapidopsis nuclei were simulated as ellipses with a major axis length of 12 1m and a minor axis length of 6 1m, and we randomly selected points using a uniform distribution in the ellipse surface.

To measure total (n_tot_), extrinsic (n_ext_) and intrinsic (n_int_) noise for cells with any number of alleles, we used an extension of the approaches previously developed ^36,37^. For a population of cells with n alleles, we considered the brightness of the transcription sites of the n alleles, expressed in number of full-length pre-mRNA molecules, as n random variables. We computed the corresponding variance-covariance matrix of these n variables and defined n_tot_ as the root square of the mean of the variances given by the matrix diagonal, n_ext_ as the root square of the mean of the covariances obtained by the non-diagonal values of the matrix, and n_int_ as the root square of n_tot_^2^-n_ext_^2^. The error in n_tot_, n_int_ and n_ext_ was estimated by randomly attributing each of the n transcription sites to the n alleles, calculating n_tot_, n_int_ and n_ext_ and repeating this 100 times to calculate the standard deviation of the values obtained.

In Figure 2D, he number of expressing cells (i.e. cells having at least 20 RNA molecules in the nucleus or the cytoplasm), as well as the number of transcription site per cell (see Fig. 4C-D), was counted manually from 2D MIP smFISH images. The plot of Figure 4 were generated in R and incorporated in the figure using Adobe Illustrator.

### Rootchips and *in vivo* live plants imaging experiments

For these experiments, we used brightest line obtained with *SPX1* promoter (the S and J lines). Transgenic seeds were germinated in conical cylinders produced from micropipette tips filled with -P agar and inserted into sterile -P agar plates. 5 to 7 day old Pi deficient seedlings were then transferred into a RootChips ^22,38^, grown vertically and fed with a Pi deficient nutritive solution at a low flow pressure until the root grew in the channel. For refeeding experiment, the nutritive solution was changed from –Pi to +Pi. Synaptored^TM^C2 (5µg/mL, Ozyme) was added into the +P solution to detect the arrival time of Pi (defining t=0). In vivo movies were recorded with an Andor Dragonfly spinning disk mounted on a Nikon Eclipse Ti using 40x Plan-Apo water objective (1.15 NA; 0.6 mm DT). Z-stacks were made every 3 minutes with a scan size of 100-150µm (0.5µm step size; ∼200 steps per Z-stack). The data was analyzed with the ImageJ software using the TrackMate plugin to track transcription sites on 2D MIPs. The intensity of *SPX1* transcription sites was assessed for several nuclei per sample. In each nucleus, the mean intensity of the nuclear background was subtracted to the maximum intensity of *SPX1* transcription site, and the resulting intensities were plotted against time.

## Acknowledgements

S.H. was supported by a PhD fellowship from the CEA and PACA region, ANR Reglisse 13-ADAP-008 fellowship and CEA DRF impulsion program with FOSSI project supported EM, LC, LN, MCT, PD. Additional grant support were also received by HJ from CEA-Enhanced Eurotalent and ANR PhlowZ 19-CE-13-0007. We thank Heliobiotech platform for the access to their RTqPCR machine. We thank Dr E. Basyuk for her help with the MS2 plasmids, Dr L. Laplaze and Dr G. Desbrosses for providing access to the growth chambers of IRD and Montpellier University. We thank Dr. O. Radulescu for his help with calculating total, extrinsic and intrinsic noise for an undefined number of alleles; Dr T. Desnos and C. Mercier for their assistance on Figures drawing and Dr S. Kanno and Dr H. Garcia for critical reading of the manuscript. We express our gratitude to the Dr J.M. Escudier for the synthesis of the SPX1 set of fluorescent probes.

We acknowledge the MRI imaging facility (belonging to the National Infrastructure France-BioImaging supported by the French National Research Agency, ANR-10-INBS-04), and the ZoOM platform (supported by the Région Provence Alpes Côte d’Azur, the Conseil General of Bouches du Rhône, the French Ministry of Research, the Centre National de la Recherche Scientifique and the Commissariat à l’Energie Atomique et aux Energies Alternatives).

## Author contributions

E.B. provided MS2 and MCP original constructs and LN conceived the experiments. L.C., S.H., P.D. performed all the experiments under LN supervision for physiological part and EB supervision for cell biology. RNA seq data were produced by D.S. and J.W. and analyzed by L.N., L.C. M-C.T. and E.M. Luminescence experiments were performed by N.P. under H.J. supervision. H.J. also implemented microfluidic technique in the SAVE team. Manuscript was written by LN and EB with help from S.H., P.D. and L.C.

## Figure Legends

**Supplemental Figure 1: Identification of genes rapidly responding to Pi resupply.**

(A) List of the fastest genes (presenting a ½Fold Change (Log_2_)½ >1) common in all the replenish RNAseq points in roots (30, 60 and 180 min), accompanied with the values obtained in shoots. Their possible regulation by PHR1/PHL1 has also been tested: presence of P1BS box 3kb upstream or 3 Kb downstream, and misregulation in *phr1/phl1* double mutant (data from Bustos et al., 2010, or analyzed by RTqPCRs and labelled* (data not shown).

(B-C-D) RTqPCR analysis of the effect of Pi replenishing for different markers WT (B-C-D) or transgenic pSQD2::LUC (C) or p35S::SPX1-GFP (D) lines. In the p35S::SPX1-GFP line, repression triggers by Pi resupply does not affect SPX1-GFP transcript whereas SPX3 mRNA (close homologue of SPX1, identified as early responding genes in the S1A list) remains strongly regulated as observed in the WT background.

The X-axis legend refers for (B) to markers analyzed in the WT background, whereas for (C and D) it refers to genetic background used (WT or transgenic), markers analyzed (LUC, GFP, SPX3) are mentioned above the graphs.

*TUBULIN* was used as housekeeping gene. Values are log_10_ relative expression to -Pi levels (REL), which are normalized to 1; n=3-5. Different letters indicate significantly different means (SNK one-way ANOVA, p<0.05, Rstudio). Error bar represent standard deviation and n the number of biological replicates used for RNA extraction.

**Supplemental Figure 2: Implementation and validation of the MS2 system in plants.**

(A) Moclo vectors adapted to the Golden Gate system and cloning strategy used to introduce the MS2 system into plants.

(B) Images are maximal image projections of microcopy images from Arabidopsis roots grown in Pi depleted medium and processed for smFISH with probes hybridizing against the endogenous *SPX1* mRNAs. Top: SmFISH signal; bottom: merge with Dapi (blue); right panels: zoom over the boxed area. Scale bars: 40 µm (left), and 4 µm (right). Pink arrows: single RNA molecules; orange arrows: transcription sites.

(C) Graph depicting the distribution of the brightness of active transcription sites for the endogenous *SPX1* gene, for cells having 1 or 2 transcription sites (yellow bars), or more than 3 (blue bars). Brightness values are expressed in number of RNA molecules (x-axis), and the y-axis represents the frequencies of these values. a.u.: arbitrary units.

**Supplemental Figure 3: Genetic and molecular analysis of selected lines (J, S) expressing the pSPX1:MS2×128 transgene.**

(A) Segregation analysis of Kanamycin resistance in the T2 progeny. Seeds were germinated on Hoagland/2 medium supplemented with kanamycin (50 mg/l). (+) = growth, (-) = death. c^2^ values are provided for theoretical segregation ratio (3:1). The critical value for a p level of 0.05 is 3.84).

(B) Molecular analysis by qPCR of T-DNA presence in the Col, S and J lines. Set of primers used are described in the Supplemental Table1 to amplify GFP, MCP and the SPX1 promoter (the only gene present in all lines). Values are presented as relative amounts (REL) normalized to S line (100%). The endogenous SPX1 gene was used as reference gene to normalize the DNA amount between samples.

**Supplemental Figure 4: The RNA produced by the SPX::MS2×128 transgene is capped and polyadenylated.**

RT-PCR amplification of the MS2×128 and endogenous SPX1 RNA after total RNA immunoprecipitation using m7G antibody (m7G-RIP). After RIP, RNAs were reverse-transcribed using oligo(dT) prior to PCR amplification. Sets of primers targeting transgene or endogenous gene were used. Wild-type (Col0) and two different transgenic lines (S and J) were used. As negative control, total RNAs were treated with pyrophosphatase to remove the cap prior to RIP experiment and RT-PCR. I : Input fraction. E : Eluate fraction. L : Ladder.

**Supplemental Figure 5: MS2 smFISH in control Col0 plants and homozygous pSPX1:MS2×128 S line grown without Pi.**

(A) Microscopy images of squashed root caps of Col0 (right two panels) and transgenic S line (left two panels), grown without Pi and processed for smFISH with probes hybridizing to the MS2×128 sequence. Top: smFISH signals; bottom: smFISH signals colored in red and merged with Dapi (blue). Images are maximal projections of z-stacks (widefield microscopy). Scale bars: 40 µm (1^st^ and 3^d^ panels), and 4 µm (2^d^ and 4^th^ panels, zooms corresponding to the boxed area of the other panels). Transcription sites and single RNA molecules are indicated by orange and pink arrows, respectively.

(B) Legend as in A, but imaging was done on mature root tissue.

**Supplemental Figure 6: SPX1 smFISH in Col0 and control spx1/spx2 mutant plants grown without Pi.**

(A) Microscopy images of squashed root caps of Col0 (left two panels) and spx1/spx2 control plants (right two panels), grown without Pi and processed for smFISH with probes hybridizing to the SPX1 mRNA. Top: smFISH signals; bottom: smFISH signals colored in red and merged with Dapi (blue). Images are maximal projections of z-stacks (widefield microscopy). Scale bars: 40 µm (1^st^ and 3^d^ panels), and 4 µm (2^d^ and 4^th^ panels, zooms corresponding to the boxed area of the other panels). Transcription sites and single RNA molecules are indicated by orange and pink arrows, respectively.

(B) Legend as in A, but imaging was done on mature root tissue.

**Supplemental Figure 7: Imaging pUnicorn1::MS2×128 and pSPX1:MS2×128 J line in fixed plant tissues.**

(A) Microscopy images of squashed mature tissue of the homozygous J line expressing pSPX1::MS2×128 grown without Pi and processed for smFISH with probes hybridizing to the MS2 sequence. Images are maximal image projections of z-stacks taken in three colors. Left (and red in the Merge panel): smFISH signals obtained with probes against the MS2 repeat; middle (and green in the Merge panel): MCP-GFP signals. Blue: nuclei stained with Dapi. The bottom panels are zooms of the boxed area in Merge panel. Orange arrows: transcription sites. Scale bars: 40 µm (top panels) and 4 µm (bottom panels).

(B) Legend as in B, except that a pUnicorn1::MS2×128 transgenic plant was imaged.

**Supplemental Figure 8: Combining microfluidics and MS2 technology reveals the fast transcriptional repression triggered by Pi supply in the S line.**

(A) Image is a maximal image projection from a time-lapse movie recorded in 3D (200 z-planes) in the pSPX1::MS2×128 S line, and displays MCP-GFP fluorescence in plants at the start of Pi refeeding (t=0; set when the Synaptored dye is detected). Scale bar: 50 µm. Inset: identical image with a higher contrast to display the tissue structure.

(B and C) Images are maximal image projection from the time-lapse movie shown in A, taken at the indicated time points after Pi refeeding and displaying MCP-GFP fluorescence. Scale bar: 10 µm.

**Supplemental Figure 9: Number of active transcription sites before and following Pi resupply in the pSPX1::MS2×128 S line.**

(A) During time lapse, the root is growing and moving in the microfluidic channel of the chip and some nuclei get out of focus. The blue curve depicts the number of nuclei still tracked at each time point (normalized to the initial number of nuclei; n_n_=113). The orange curve indicated the number of active transcription sites detected in those nuclei, normalized to number of transcription sites present when the recording started (n_t_=107).

(B) Biological replicates of (A) with n_n_= 62, 66, 94, 81 and n_t_= 49, 72, 45, 47 for plants 2 to 5 respectively.

**Supplemental Figure 10: Transcription site activity following Pi resupply in the pSPX1::MS2×128 J line transgenic plants.**

(A) Normalized intensities of fluorescence signals at transcription sites over time, recorded every 3 minutes after Pi refeeding, for cells exhibiting one or two active transcription sites (n=16). Normalization is done according to the value measured at t=0.

(B) Average normalized intensities of transcription sites, for the 16 transcription sites shown in panel A (mean and standard deviation).

(C) Normalized intensities of fluorescence signals at transcription sites over time, and recorded every 3 minutes after Pi refeeding, for cells exhibiting one or two active transcription sites (n=13) in an independent plant than in (A). Normalization is done according to the value measured at t=0

(D) Average normalized intensities of transcription sites, for the 13 transcription sites shown in panel B (mean and standard deviation).

**Supplemental Figure 11: Transcription site activity upon continuous exposure to Pi depleted medium in pSPX1::MS2×128 S line transgenic plants.**

(A) Maximal image projection from a time-lapse movie recorded in 3D (200 z-planes) and displaying MCP-GFP fluorescence in S plants in steady-state conditions (-Pi condition) from a mature root area. Scale bar: 100 µm. Inset: identical image with a higher contrast to display the tissue structure.

(B) The panels are zooms of the boxed area form (A) recorded in the time-lapse movie taken at the indicated time points and displaying MCP-GFP fluorescence. Scale bar: 20 µm.

**Supplemental Figure 12: Location of smFISH oligonucleotide probes on their respective targets RNA**

(A) SPX1 mRNA.

(B) MS2×128 RNA.

(C) Post polyA region of the pSPX1::MS2×128 reporter gene.

**Supplementary Table 1: Primer sequences used for RTqPCR and PCR reaction. Supplementary Table 2: Primer sequences for smFISH experiment.**

(A) Primers used to detect *SPX1* mRNA. Cy3 fluorescent dye is grafted on amino-modified uridine analogue C6dT (labelled X on the sequence).

(B) Primers used in smiFISH to detect the post polyA region of the pSPX1::MS2×128 transcripts.

(C) Primers used to detect MS2×128 transcripts.

**Movie 1 and 2: Bursting activity of the pSPX1::MS2×128 reporter**

Movie of root cap cells of Arabidopsis S line expressing pSPX1::MS2×128 and MCP-GFP, and continuously grown without Pi. Maximal image projection (XY and XZ) are from a time-lapse movie recorded in 3D (44 z-planes). Time (in min) is indicated. Movie 1 illustrates gene bursting while Movie 2 illustrates the release of single RNAs in the nucleoplasm when promoter activity stochastically turns off when a burst ends.

**Movies 3 to 8: Transcriptional repression of the pSPX1::MS2×128 reporter triggered by Pi resupply.**

Movies of root cells of Arabidopsis S (3 to 7) and J (8) line transformed with pSPX1::MS2×128 and MCP-GFP, after receiving a Pi rich solution at time t=0 min. Maximal image projection from a time-lapse movie recorded in 3D (200 z-planes). Acquisitions lasted 39 min for Movies 3 and Movie 4 (Movie 4 zooms on few cells cropped from Movie 3), 54 min for Movie 5 to Movie 7 (Movies 6 and 7 are magnifications deriving from Movie 5), and 45 min for Movie 8.

