## Supplemental Figures for "RNA labelling in live plants reveals single cell transcriptional dynamics: application to phosphate signaling"

A

| Gene ID | Gene onthology | P1BS 3kb up | P1BS 3kb down | Altered in phr1/phl1 | ROOTS +P 180 min **FKBP** | ROOTS -P 180 min **FKBP** | ROOTS -P vs +P 180 min **Log2FC** | ROOTS -P vs re 30 min **Log2FC** | ROOTS -P vs re 60 min **Log2FC** | ROOTS -P vs re 180 min **Log2FC** | LEAVES -P vs +P 180 min **FKBP** | LEAVES -P vs re 30 min **Log2FC** | LEAVES -P vs re 60 min **Log2FC** | LEAVES -P vs re 180 min **Log2FC** |
|---|---|---|---|---|---|---|---|---|---|---|---|---|---|---|
| AT5G20150 | SPX domain gene 1 (SPX1) | 3 | 2 | Yes* | 20,9 | 1566,6 | -6,23 | -1,05 | -2,25 | -2,13 | -5,35 | -0,02 | -1,18 | -2,06 |
| AT5G20790 | unknown protein (Unicorn1) | 1 | 0 | Yes | 3,0 | 250,7 | -6,36 | -1,77 | -2,09 | -2,43 | -6,74 | 0,46 | -1,70 | -3,37 |
| AT3G02040 | glycerophosphoryl diester phosphodiesterase | 4 | 0 | Yes | 6,4 | 244,0 | -5,25 | -1,21 | -2,56 | -2,15 | -5,87 | -0,14 | -1,66 | -1,00 |
| AT3G47420 | putative glycerol 3 phosphate transporter | 0 | 3 | Yes | 12,3 | 232,5 | -4,24 | -1,33 | -2,15 | -1,22 | -3,08 | 0,30 | -1,39 | -2,00 |
| AT1G08650 | phosphoenolpyruvate carboxylase kinase 1 | 3 | 2 | Yes | 36,3 | 183,8 | -2,34 | -1,97 | -1,84 | -1,41 | -2,81 | 0,22 | -1,70 | -1,44 |
| AT2G45130 | SPX domain gene 3 (SPX3) | 2 | 1 | Yes | 0,2 | 151,4 | -9,56 | -1,16 | -2,39 | -2,37 | -11,05 | 0,31 | -1,30 | -0,96 |
| AT3G30725 | glutamine dumper 6 | 0 | 1 | Yes* | 1,7 | 125,4 | -6,17 | -1,50 | -1,71 | -2,23 | -4,6 | -0,10 | -1,34 | -2,39 |
| AT1G19200 | cyclin-dependent kinase | 3 | 1 | Yes | 3,8 | 59,9 | -3,97 | -1,14 | -2,04 | -1,98 | -4,72 | 0,11 | -0,95 | -4,61 |
| AT3G07350 | sulfate/thiosulfate import ATP-binding protein | 1 | 4 | Yes | 11,4 | 41,6 | -1,86 | -1,22 | -2,31 | -1,52 | -0,7 | -1,35 | -3,05 | -0,65 |
| AT3G09400 | pol-like 3 | 1 | 0 | Yes | 1,1 | 31,7 | -4,82 | -1,14 | -2,51 | -2,13 | -4,14 | -0,79 | -1,56 | -3,95 |
| AT1G72070 | chaperone DnaJ-domain superfamily protein | 2 | 1 | Yes | 4,5 | 27,0 | 2,59 | -1,11 | -1,61 | -1,69 | -1,98 | -0,38 | -0,35 | -1,71 |
| AT4G32480 | sugar phosphate exchanger | 5 | 1 | Yes* | 5,7 | 21,6 | -1,92 | -1,02 | -1,77 | -1,20 | -2,29 | -1,16 | -3,29 | -1,59 |
| AT3G26510 | octicosapeptide/Phox/Bem1p family protein | 1 | 0 | Yes | 9,5 | 19,9 | -1,06 | -1,64 | -1,28 | -1,43 | -0,49 | -0,90 | -1,63 | -1,18 |
| AT1G71130 | ethylene response factor | 1 | 0 | Yes | 3,9 | 16,5 | -2,10 | -1,51 | -1,87 | -1,09 | -2,09 | 0,22 | -1,79 | -0,26 |
| AT5G67430 | acyl-CoA N-acyltransferases (NAT) superfamily protein | 1 | 1 | Yes | 2,7 | 14,9 | -2,44 | -1,85 | -1,41 | -1,08 | -2,35 | -1,09 | -0,49 | -1,05 |
| AT2G46940 | unknown protein | 1 | 2 | Yes | 3,1 | 10,6 | -1,78 | -1,34 | -1,59 | -1,45 | -2,72 | 0,96 | 0,05 | -2,11 |
| AT3G29670 | HXXXD-type acyl-transferase family protein | 1 | 1 | Yes | 0,8 | 10,6 | -3,70 | -1,74 | -1,42 | -1,15 | -0,46 | -2,05 | -1,72 | -1,70 |
| AT4G30370 | IRT1 degradation factor1 | 1 | 1 | Yes* | 2,6 | 8,9 | -1,80 | -1,02 | -1,33 | -1,70 | -2,38 | 0,16 | -1,74 | -2,54 |
| AT1G54095 | unknown protein (DUF1677) | 1 | 0 | Yes* | 3,3 | 8,9 | -1,41 | -1,25 | -1,67 | -1,29 | -1,45 | -1,18 | 0,22 | -3,61 |
| AT2G07042 | other RNA | 1 | 0 | No* | 0,5 | 4,7 | -3,15 | -1,85 | -2,03 | -2,07 | -0,41 | 0,20 | -2,12 | -3,20 |
| AT2G26150 | heat shock transcription factor A2 | 0 | 0 | No | 1,3 | 4,6 | -1,83 | -4,58 | -1,80 | -1,65 | -2,74 | -3,09 | -1,66 | -3,34 |
| AT5G09470 | dicarboxylate carrier 3 | 0 | 2 | Yes | 0,1 | 1,4 | -3,62 | -1,18 | -1,77 | -2,01 | -7,77 | 0,37 | -1,04 | -0,41 |

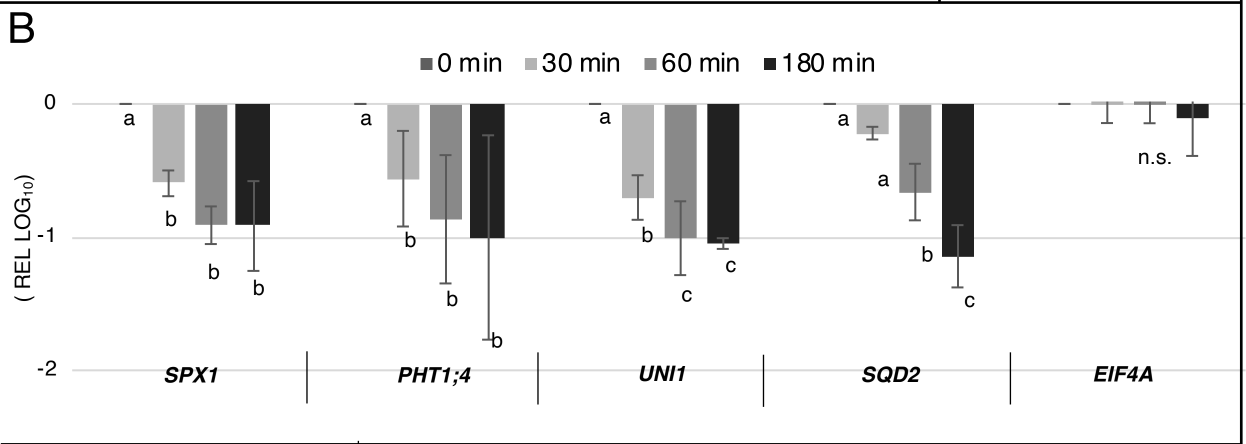


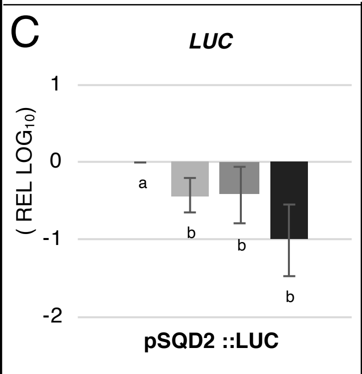


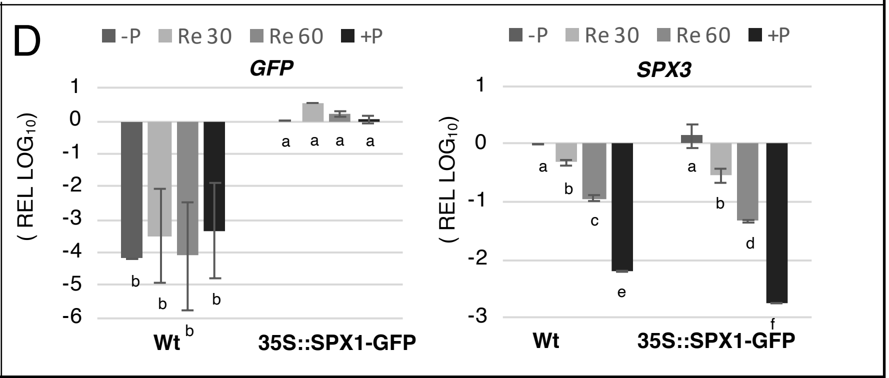


Figure S1

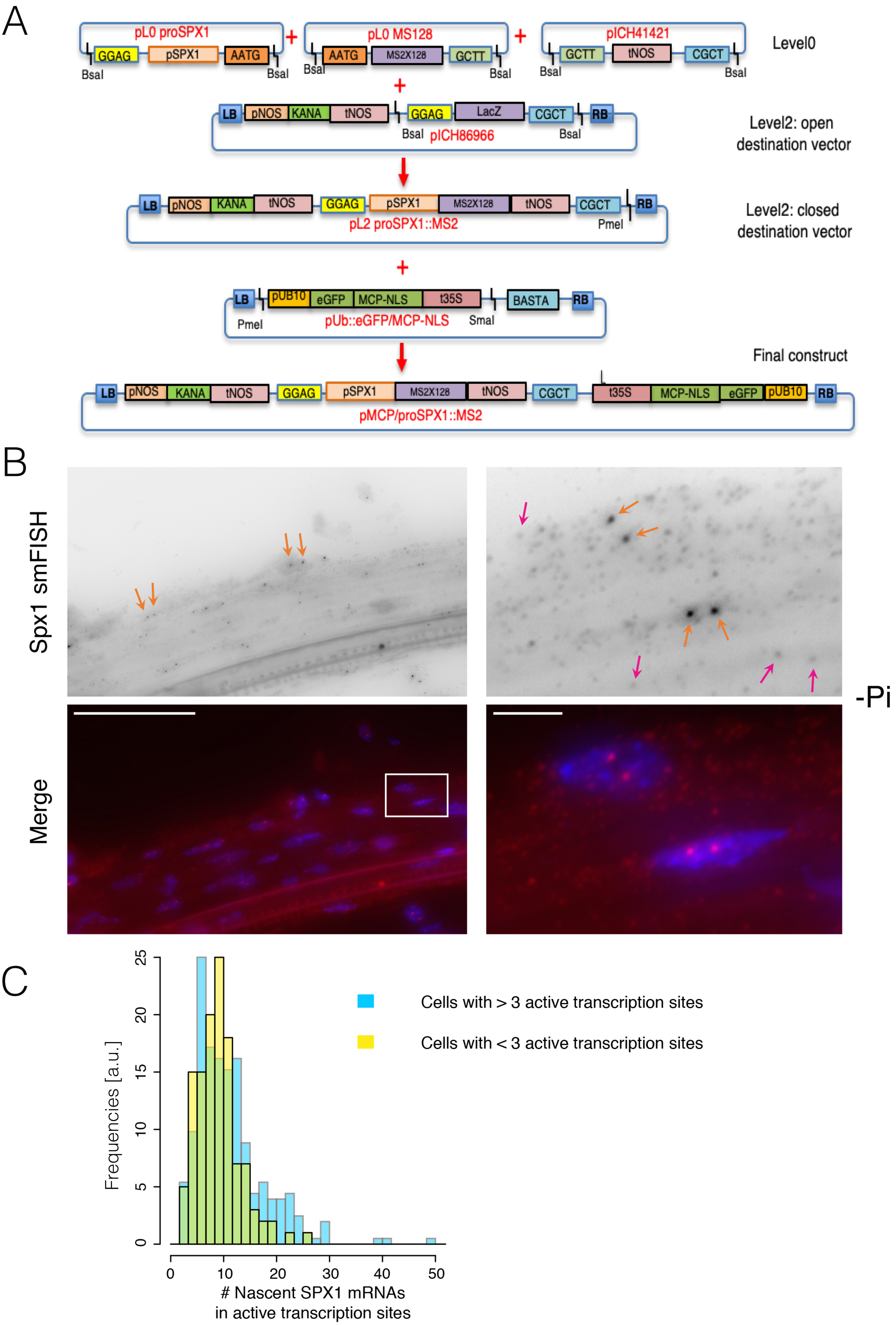
A
pL0 proSPX1
GGAG
pSPX1
AATG
BsaI
BsaI
pL0 MS128
AATG
MS2X128
GCTT
BsaI
BsaI
pICH41421
GCTT
tNOS
CGCT
BsaI
BsaI
Level0
LB
pNOS
KANA
tNOS
GGAG
LacZ
CGCT
RB
BsaI
pICH86966
BsaI
Level2: open destination vector
LB
pNOS
KANA
tNOS
GGAG
pSPX1
MS2X128
tNOS
CGCT
RB
pL2 proSPX1::MS2
PmeI
Level2: closed destination vector
LB
pUB10
eGFP
MCP-NLS
t35S
BASTA
RB
PmeI
pUb::eGFP/MCP-NLS
SmaI
Final construct
LB
pNOS
KANA
tNOS
GGAG
pSPX1
MS2X128
tNOS
CGCT
t35S
MCP-NLS
eGFP
pUB10
RB
pMCP/proSPX1::MS2
B
Spx1 smFISH
Merge
-Pi
C
Frequencies [a.u.]
### Nascent SPX1 mRNAs in active transcription sites
Cells with > 3 active transcription sites
Cells with < 3 active transcription sites
0
5
10
15
20
25
0
10
20
30
40
50

Figure S2

A

| Lines | + | - | R | $\chi^2$ | Loci |
|---|---|---|---|---|---|
| S | 65 | 23 | 2.8 | 0.07 | 1 |
| J | 37 | 13 | 2.8 | 0.03 | 1 |

B

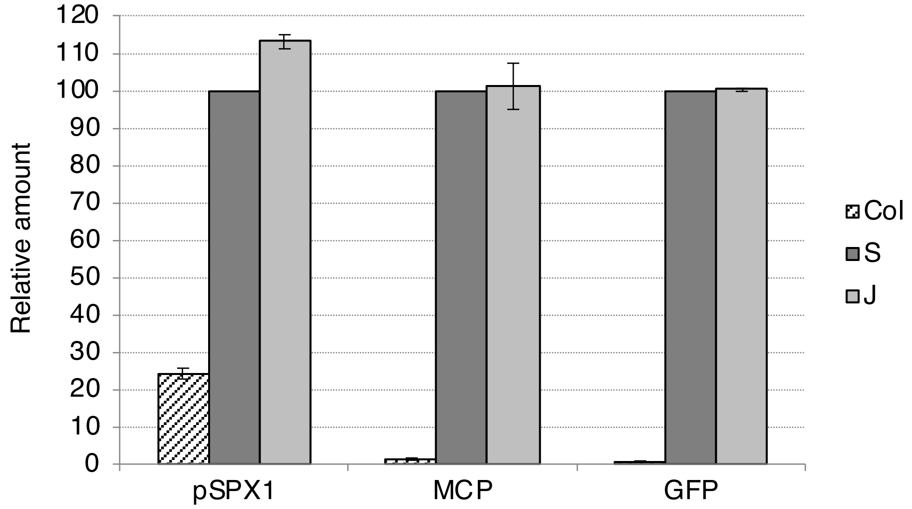


Figure S3

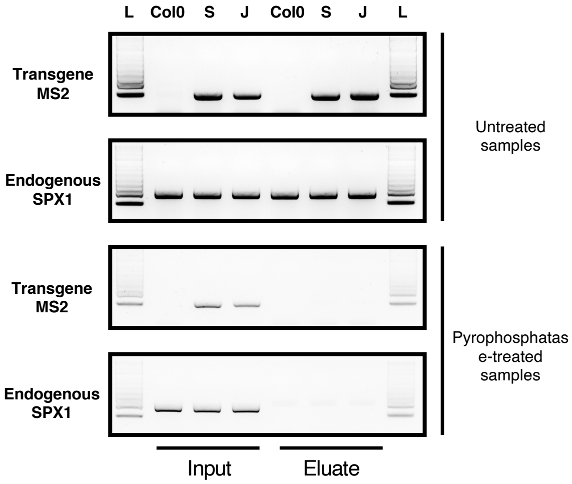
L
Col0
S
J
Col0
S
J
L
Transgene MS2
Endogenous SPX1
Transgene MS2
Endogenous SPX1
Untreated samples
Pyrophosphatase-treated samples
Input
Eluate

Figure S4

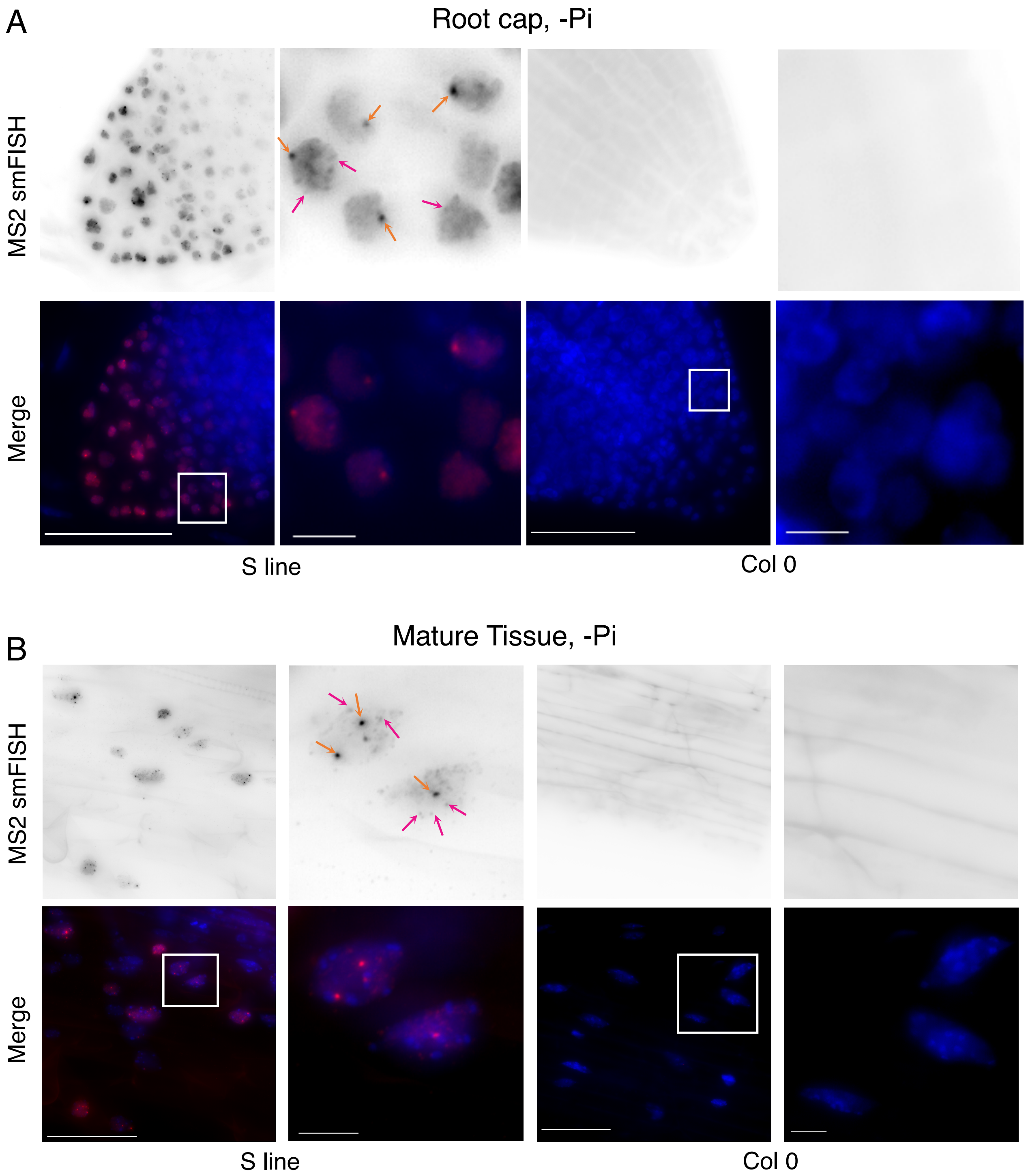
A
Root cap, -Pi
MS2 smFISH
Merge
S line
Col 0
B
Mature Tissue, -Pi
MS2 smFISH
Merge
S line
Col 0

### Figure S5

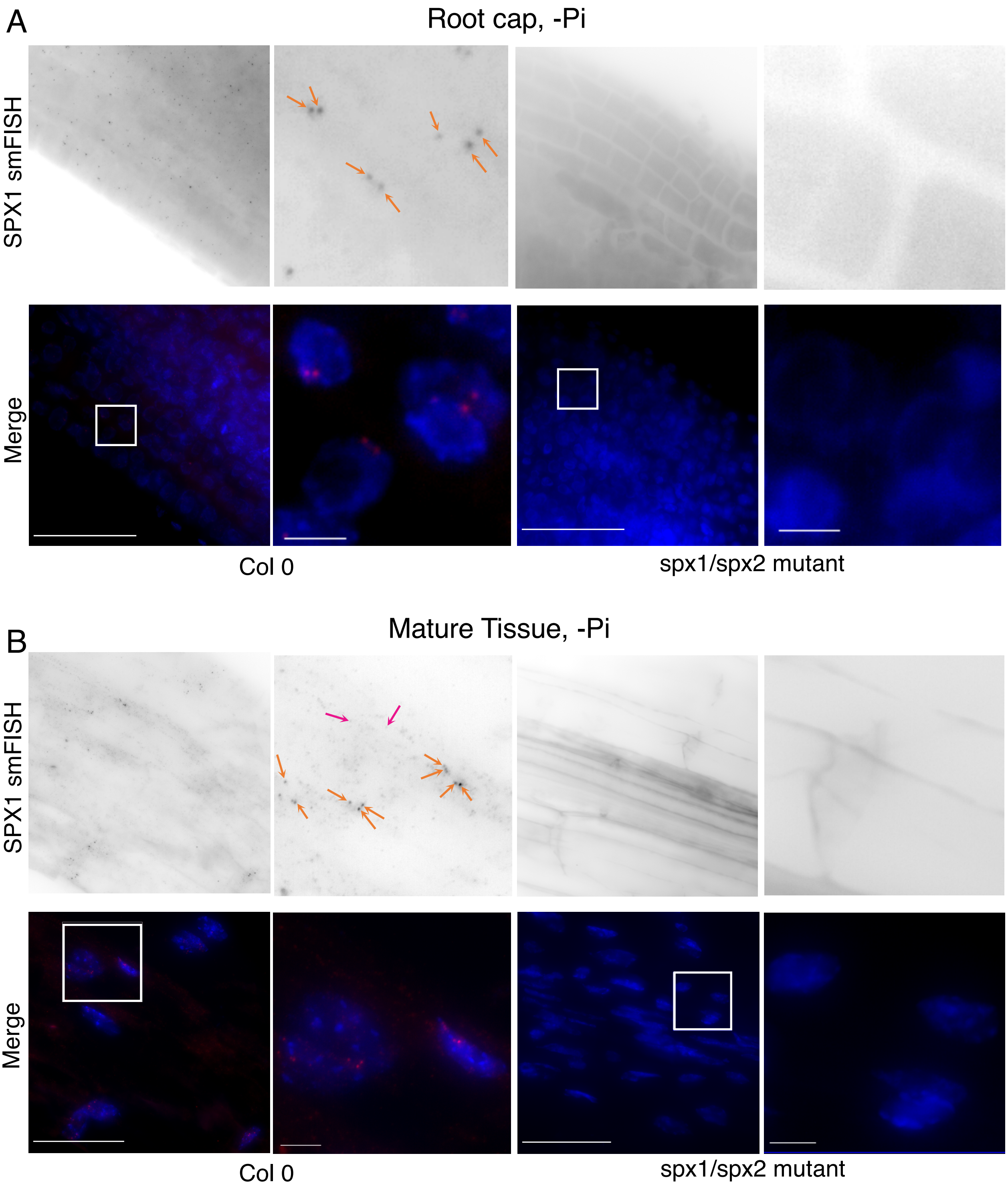
A
Root cap, -Pi
SPX1 smFISH
Merge
Col 0
spx1/spx2 mutant
B
Mature Tissue, -Pi
SPX1 smFISH
Merge
Col 0
spx1/spx2 mutant

Figure S6

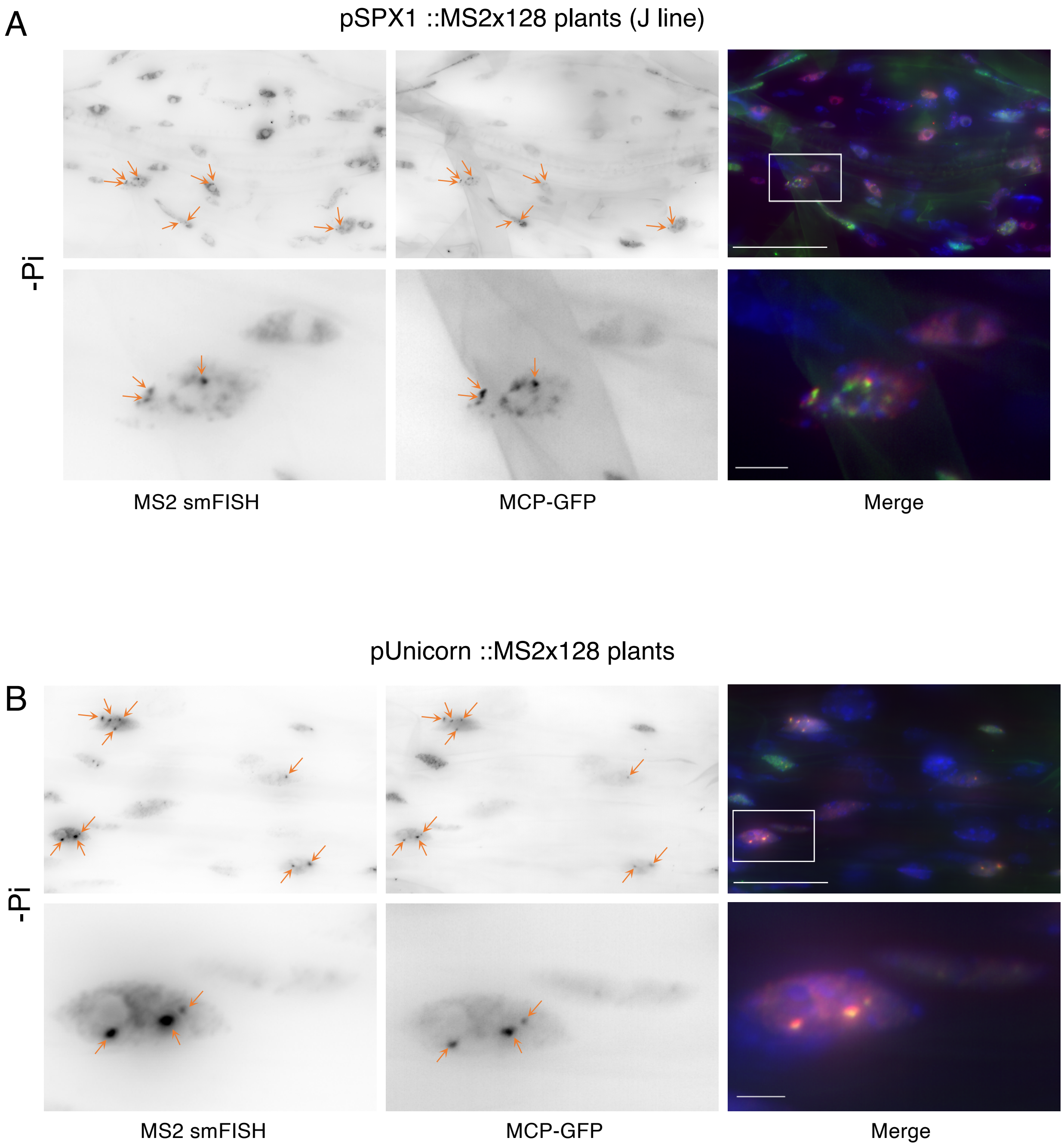
A
pSPX1 ::MS2x128 plants (J line)
-Pi
MS2 smFISH
MCP-GFP
Merge
pUnicorn ::MS2x128 plants
B
-Pi
MS2 smFISH
MCP-GFP
Merge

Figure S7

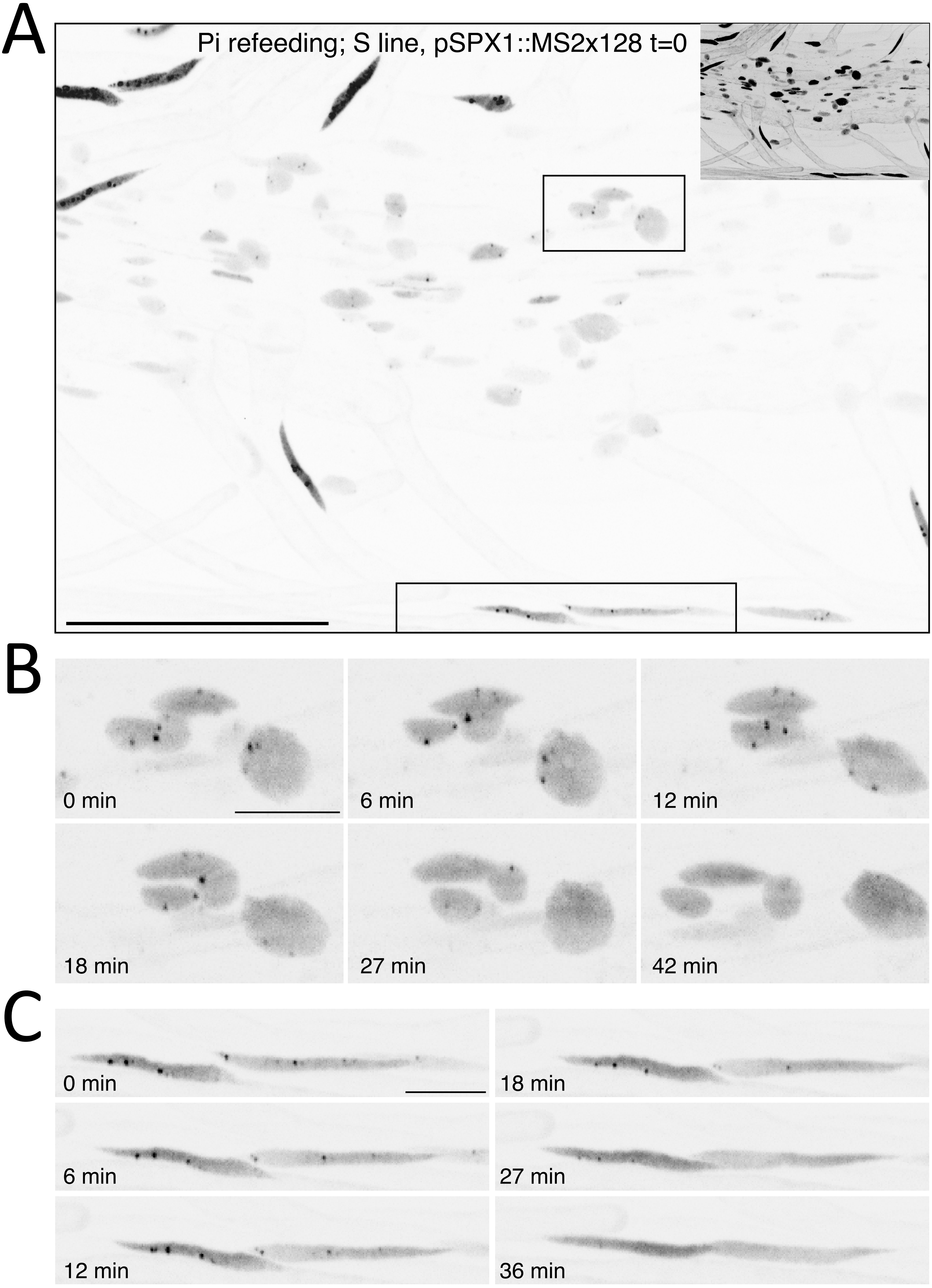
A
Pi refeeding; S line, pSPX1::MS2x128 t=0
B
0 min
6 min
12 min
18 min
27 min
42 min
C
0 min
18 min
6 min
27 min
12 min
36 min

Figure S4

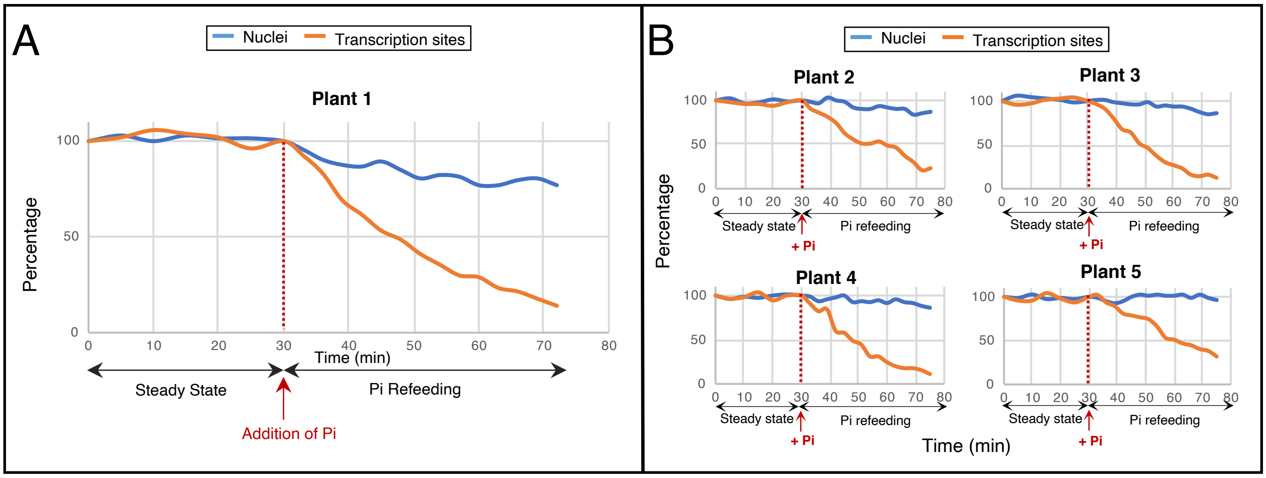
A
Nuclei
Transcription sites
Plant 1
Percentage
Time (min)
Steady State
Pi Refeeding
Addition of Pi
B
Plant 2
Plant 3
Plant 4
Plant 5
Steady state
Pi refeeding
+ Pi

Figure S9

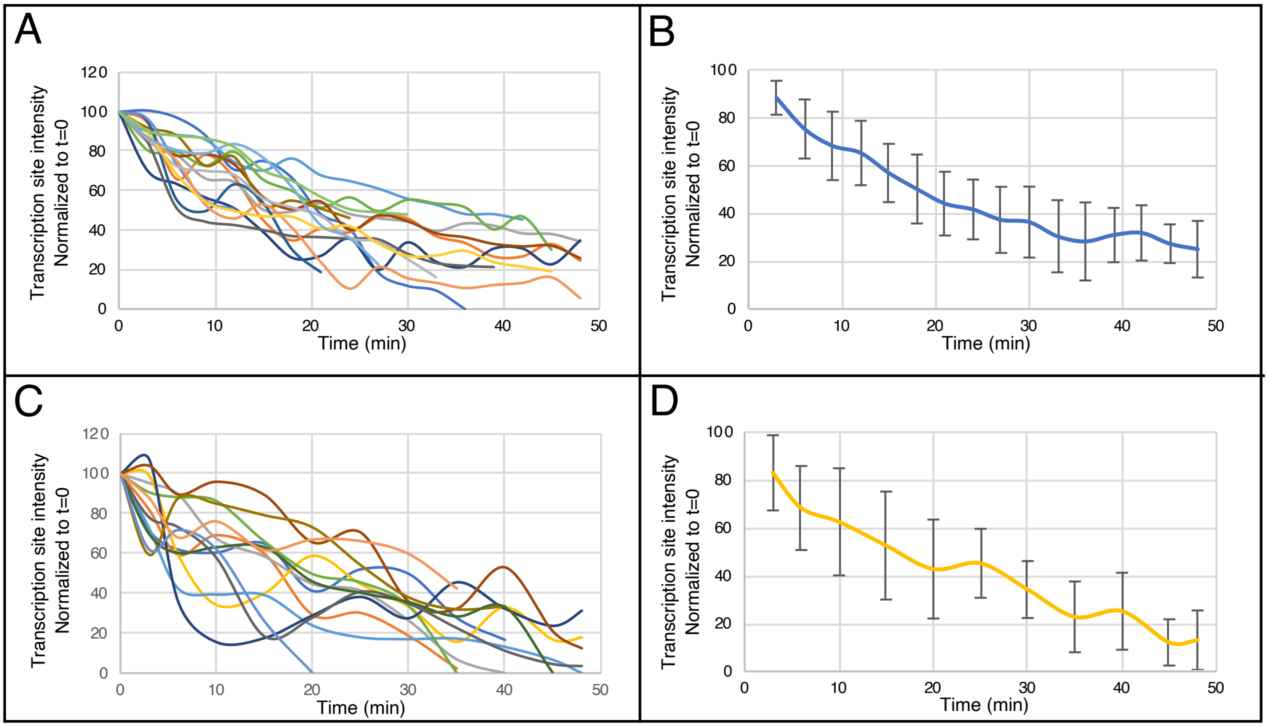
A
B
C
D
Transcription site intensity
Normalized to t=0
Time (min)
120
100
80
60
40
20
0
10
20
30
40
50

Figure S10

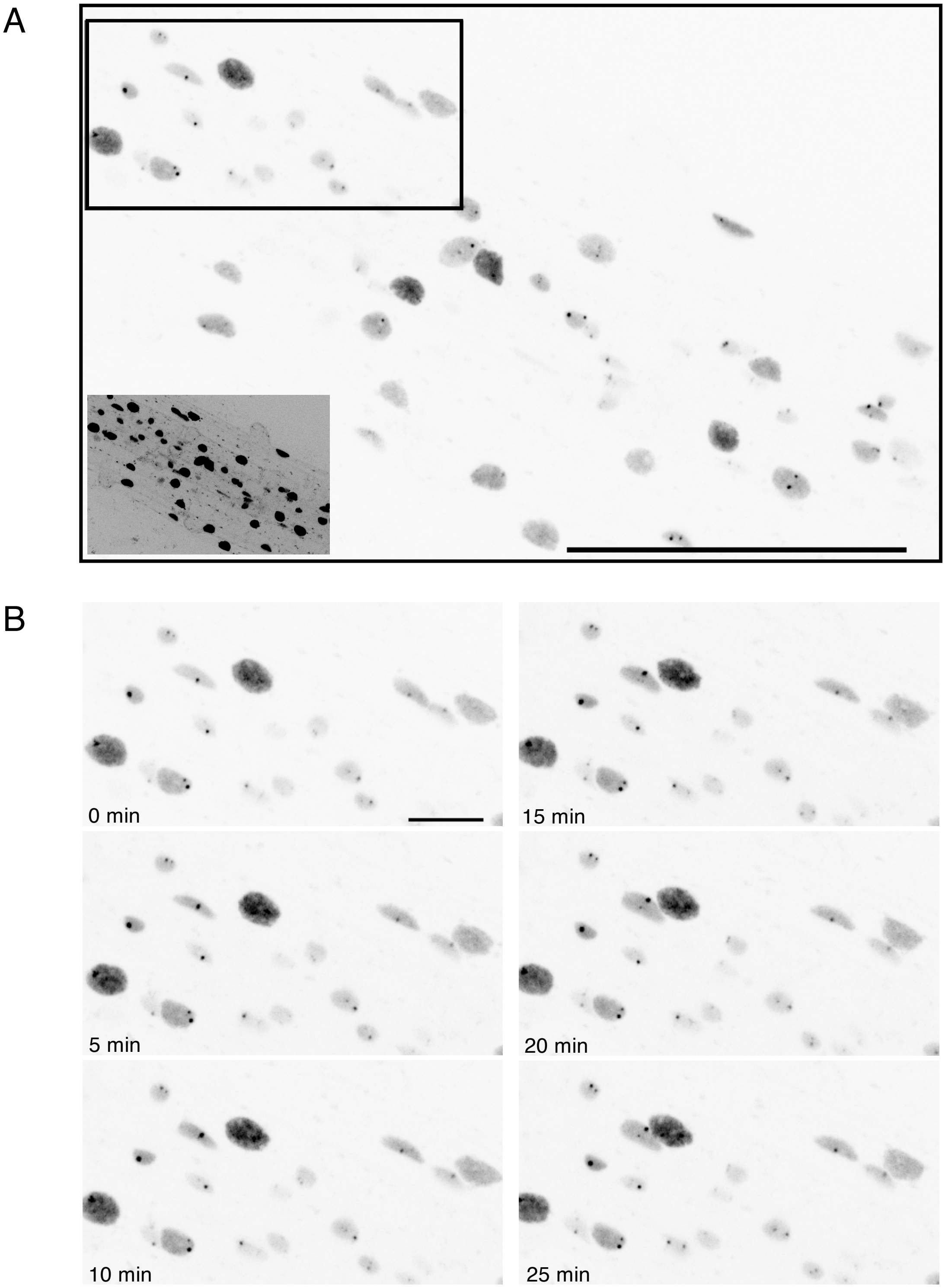


Figure S11

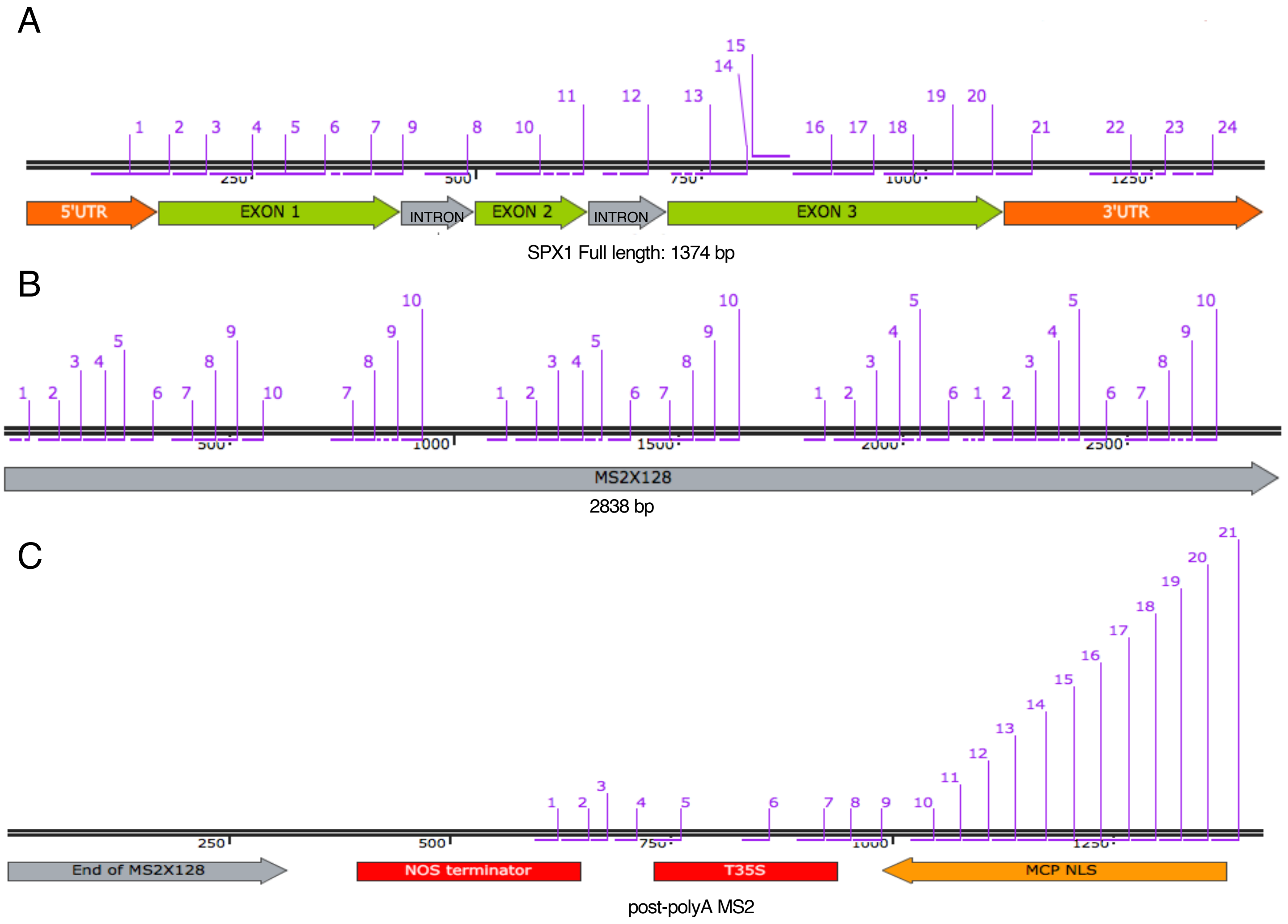
A
5'UTR
EXON 1
INTRON
EXON 2
INTRON
EXON 3
3'UTR
SPX1 Full length: 1374 bp
B
MS2X128
2838 bp
C
End of MS2X128
NOS terminator
T35S
MCP NLS
post-polyA MS2

Figure S12
